# Strain-level metagenomic profiling using pangenome graphs with PanTax

**DOI:** 10.1101/2025.04.29.651271

**Authors:** Wenhai Zhang, Yuansheng Liu, Guangyi Li, Jialu Xu, Enlian Chen, Alexander Schönhuth, Xiao Luo

## Abstract

Microbes are omnipresent, thriving in a range of habitats, from oceans to soils, and even within our gastrointestinal tracts. They play a vital role in maintaining ecological equilibrium and promoting the health of their hosts. Consequently, understanding the diversity in terms of strains in microbial communities is crucial, as variations between strains can lead to different phenotypic expressions or diverse biological functions. However, current methods for taxonomic classification from metagenomic sequencing data have several limitations, including their reliance solely on species resolution, support for either short or long reads, or their confinement to a given single species. Most notably, most existing strain-level taxonomic classifiers rely on the sequence representation of multiple linear reference genomes, which fails to capture the sequence correlations among these genomes, potentially introducing ambiguity and biases in metagenomic profiling.

Here, we present PanTax, a pangenome graph-based taxonomic profiler that overcomes the short-comings of sequence-based approaches, because pangenome graphs possess the capability to depict the full range of genetic variability present across multiple evolutionarily or environmentally related genomes. PanTax provides a comprehensive solution to taxonomic classification for strain resolution, compatibility with both short and long reads, and compatibility with single or multiple species. Extensive benchmarking results demonstrate that PanTax drastically outperforms state-of-the-art approaches, primarily evidenced by its significantly higher precision or recall at strain level, while maintaining comparable or better performance in other aspects across various datasets. PanTax is an open-source user-friendly tool that is publicly available at https://github.com/LuoGroup2023/PanTax.

## Introduction

Microorganisms are ubiquitous on Earth, inhabiting diverse environments such as oceans, soils, and gastrointestinal tracts, where they play indispensable roles in maintaining ecological balance and host health. Microbial communities are often composed of multiple types of microorganisms, at uneven species abundance and high diversity. Rapid mutation and horizontal gene transfer in microorganisms may result in different strains of the same species, which further increases the biological diversity and complexity of microbial communities. Diversity of microbial communities at the level of strains is a key factor in microbiome-related research, as it does not only reflect the evolution and adaptation of microorganisms, but also their interactions with the environment and the functions of their host. Previous studies have shown that different strains may exhibit different phenotypes or perform different biological functions in the environment (Marx, 2016; Van Rossum *et al*., 2020). Moreover, recent works have revealed that strain-level genome variations in the gut microbiota are closely related to host health and disease (Zhernakova *et al*., 2024; Chen *et al*., 2022; Zeevi *et al*., 2019). This means that determining the composition of gut microbiota in terms of their strains can accurately predict intestinal diseases such as inflammatory bowel disease (Jiang *et al*., 2022), and is important for understanding the biological characteristics of microorganisms in general and their involvement in scenarios of clinical interest in particular (Bickhart *et al*., 2022; Zheng *et al*., 2022; Fedarko *et al*., 2022). Therefore, to reveal the structure and function of microbial communities, we need to estimate their composition and abundance at finest resolution possible, that is, not only at the level of species, but also at the level of strains.

For this study, we define a “strain” as a microbial genome with sufficient genomic variation to be distinguished from other genomes within the same species. This definition aligns with previous studies (Vicedomini *et al*., 2021; Liao *et al*., 2023), although we acknowledge that the term “strain” may be interpreted differently in traditional taxonomy and other areas of microbial evolution.

Metagenome sequencing preserves the majority of the genetic information and enables a strain-level analysis of microbial communities. This establishes significant advantages over amplicon sequencing and culture-based methods, because these two approaches, by design, inevitably miss large parts of the genetic material one seeks to analyze. Next generation sequencing (NGS) has been routinely used in reference-based taxonomic classification of metagenomic data, which explains the development of many related tools. The fundamental tasks are to classify or group sequencing reads into distinct genomic bins, each representing a different taxon (e.g., species or strain) within a metagenomic sample *(taxonomic binning)*, and to identify the taxa present while estimating their relative abundances in the sample *(taxonomic profiling)* (Simon *et al*., 2019; Sun *et al*., 2021). Despite their differences, these tools are frequently used interchangeably in the metagenomic data analysis.

Various tools have been developed that operate at the level of species alone. However, strain-level analysis has still remained a tough challenge. All of the species-level tools receive the sequenced reads of a metagenome as input, and output the spectrum of species that make part of the metagenome, and also possibly their relative sequence abundances (the proportion of reads assigned to a given taxon relative to the total reads) or relative taxonomic abundances (the proportion of genomes of a given taxon to the total genomes detected) (Simon *et al*., 2019). Note that none of these tools aims to assemble the individual genomes prior to raising the relevant estimates. Rather, they put the reads of a metagenome into context with existing reference sequences and derive the corresponding conclusions from the resulting read-to-reference sequence alignments.

Existing tools can be categorized into three types. The first are marker-based methods, such as MetaPhlAn2 (Truong *et al*., 2015), MetaPhlAn4 (Blanco-Míguez *et al*., 2023), and mOTUs2 (Milanese *et al*., 2019). These methods utilize a subset of marker genes, typically gene families that are indicative of species. For example, while the 16S rRNA sequence is commonly used in some microbiome analysis methods due to its high conservation (Edgar, 2018), tools like MetaPhlAn and mOTUs rely on a broader set of specific taxon-specific markers. While these methods are computationally efficient, the classification can be biased if the marker genes in the microbial sequences of interest do not follow a uniform distribution (D’Amore *et al*., 2016).

The second are DNA-to-protein methods, where DIAMOND (Buchfink *et al*., 2015), Kaiju (Menzel *et al*., 2016) and MMseqs2 (Steinegger and Söding, 2017) are prominent examples. DNA-to-protein methods focus on the protein-coding sequences of the genomes, while discarding all non-coding sequence from further consideration. This prevents the monitoring of differences that occur in the non-coding portions of the bacterial genomes. The third are DNA-to-DNA methods, such as Kraken (Wood and Salzberg, 2014), Kraken2 (Wood *et al*., 2019), KrakenUniq (Breitwieser *et al*., 2018), Bracken (Lu *et al*., 2017), CLARK (Ounit *et al*., 2015), Centrifuge (Kim *et al*., 2016), Centrifuger (Song and Langmead, 2024), CAMMiQ (Zhu *et al*., 2022), KMCP (Shen *et al*., 2023), Ganon (Piro and Reinert, 2023) and Sylph (Shaw and Yu, 2024) characterized by considering whole-genome information for classification.

However, most of these methods were originally designed for taxonomic classification at the species level and do not perform as effectively at the strain level. Strain-level microbiome composition analysis tools that have been suggested recently, such as StrainScan (Liao *et al*., 2023), StrainGE (van Dijk *et al*., 2022) or StrainEst (Albanese and Donati, 2017), can only handle one species/genus specified by the user as input. They can only handle one species/genus that preassigned by users. These tools are specifically designed to address the unique characteristics of metagenomes, distinguishing them from tools used for classifying isolated genomes, such as those derived from individual organisms grown on Petri dishes or from environments that do not contain whole microbial communities. Approaches that address the classification of isolates, but not metagenomes, are ProPhyle (Břinda *et al*., 2017) and Phylign (Břinda *et al*., 2023), for example.

NGS or short-read sequencing, characterized by reads of a few hundred base pairs (bp) in length, has been the preferred method for metagenomics due to its ubiquitous availability and its low cost. In the meantime, third generation sequencing (TGS) or long-read sequencing technologies such as Pacific Biosciences (PacBio) and Oxford Nanopore Technologies (ONT), have become more affordable. TGS has three advantages over short-read sequencing when dealing with metagenomic assignment and estimating the bacterial composition. First, long reads, of length tens of Kbp to Mbp, retain much longer-range genomic information. Therefore, they can more easily distinguish intra-genomic repeats or highly similar strains, which supports the disambiguation of reads when determining the strain they stem from. Second, as per the basic properties of single-molecule sequencing technologies, TGS (PacBio and ONT) avoid the generation of PCR duplicates common to NGS, which avoids the corresponding biases during taxonomic profiling. Third, portable TGS sequencers, such as ONT Flongle and MinION, enable cost-effective, in-field and real-time metagenomic sequencing. This can be used in scenarios where speed of identification of microbial/viral components matters, such as, for example, during pandemics (Quick *et al*., 2016; Brinkmann *et al*., 2021). Urgency may also be a factor in scenarios where samples are difficult to culture or preserve (Runtuwene *et al*., 2019; Wang *et al*., 2021), or when analyzing samples raised in hospitals (Chng *et al*., 2020).

Still, the largest part of taxonomic classifiers is based on NGS. Since the corresponding built-in algorithms are not suitable for querying sequences that are long and/or affected by high error rates, their underlying technology cannot be straightforwardly adapted to processing TGS reads. Therefore, several NGS-oriented tools, such as Kraken2, KrakenUniq, KMCP, and Ganon, can successfully process TGS data. However, they may encounter challenges in yielding satisfactory results.

The difficulties involved in the adaptation of tools from NGS to TGS, as just outlined, explains why developing novel approaches for TGS-based taxonomic classification is imperative. So far, only few related tools have been designed, such as Melon (marker-based) (Chen *et al*., 2023), MEGAN-LR (DNA-to-protein) (Huson *et al*., 2018) and MetaMaps (DNA-to-DNA) (Dilthey *et al*., 2019). Just as for the tools already listed above, the characteristic drawbacks of the above-mentioned three categories apply also here. When raising a summarizing overall account of the characteristics of tools (see Table 1 for an overview), Centrifuge (Kim *et al*., 2016) and Centrifuger (Song and Langmead, 2024) emerges as the methods that can handle both NGS and TGS reads and perform strain-level taxonomic profiling for multiple species at a time, so proves to be the tools already available to address all currently driving issues in a universal manner.

**Table 1.**
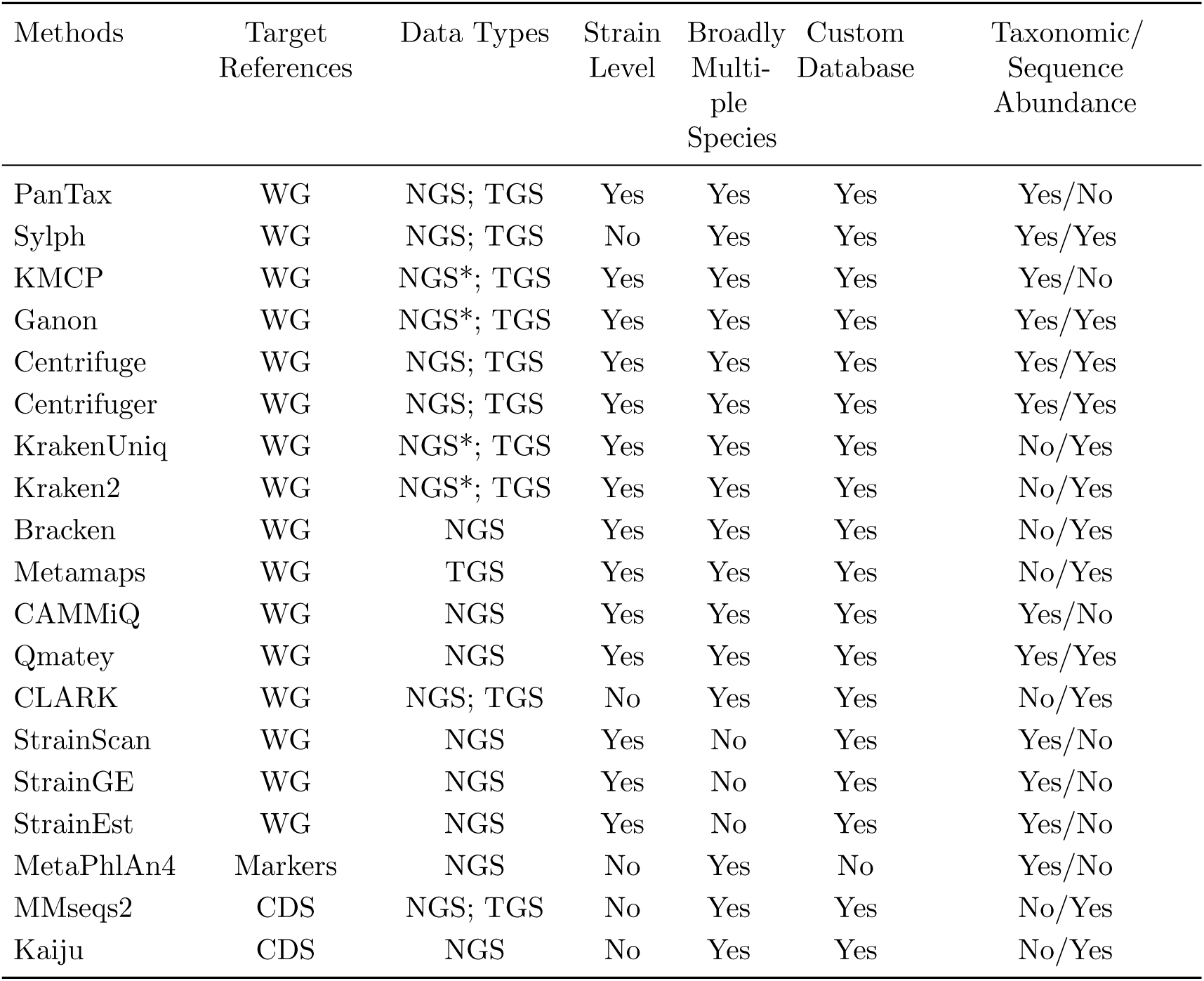
Characteristic summary of representative taxonomic classifiers. WG: whole genome. CDS: coding sequence. * indicates that the tool was primarily developed and tested for NGS data but can also be successfully ran on TGS data.

To overcome the shortcomings of existing taxonomic metagenome classifiers, we suggest PanTax ([Pan]genome graph based [Tax]onomic classifier), as a new method to perform strain-level taxonomic profiling for both short-read and long-read metagenomic data. From a broader perspective, PanTax refers to the pangenome that captures the genomes of multiple strains as a reference, instead of only a single reference genome for each species. The corresponding expansion in terms of strain diversity that PanTax builds on enhances both the accuracy in mapping metagenome reads and the subsequent estimation of its composition at the level of strains. From a methodological point of view, PanTax’s innovation is the systematic integration of pangenome graphs, instead of plain linear sequences, as an algorithmic foundation for taxonomic classification. The particular type of pangenome graphs that PanTax relies on are variation graphs, as originally described in (Paten *et al*., 2017). In the meantime, variation graphs have proven effective in addressing a wide range of computational genomics challenges.

Examples are primary read mapping and variant calling (Garrison *et al*., 2018; Martiniano *et al*., 2020; Sirén *et al*., 2021), haplotype modeling (Rosen *et al*., 2017), as well as the refined correction of errors in long reads (Luo *et al*., 2022b). For the latter, greatest improvements were observed for long reads stemming from metagenomes. The favorable usage of variation graphs in the assembly of genomes from mixed samples (Baaijens *et al*., 2019, 2020) is an additional hint to the particular advantages of variation graphs in the analysis of metagenomes. To the best of our knowledge, PanTax is the first approach that employs pangenome graphs as whole genome-wide reference systems for the taxonomic profiling of metagenomes.

To provide evidence of PanTax’s superiority, we have conducted various challenging experiments and compared PanTax with the state-of-the-art taxonomic classifiers on both simulated and real datasets that relate to questions of actual interest in research. The corresponding benchmarking experiments consistently demonstrate that PanTax achieves the best performance rates in taxonomic profiling, across all popular sequencing platforms.

## Results

We have developed and implemented PanTax, a novel method for classifying the contents of metagenomes. PanTax is universal insofar as it 1) evaluates whole genomes, 2) can take both NGS or TGS reads as input, 3) can process multiple species simultaneously, 4) refers to a custom database, which avoids the biases of external databases, 5) provides estimates on the taxonomic abundances of the involved strains, As was pointed out on plenty of occasions, all of these points are of critical relevance in the analysis of metagenomes.

The foundation for PanTax’s superiority is to employ pangenome graphs, instead of linear genome sequences as a basis for the required reference systems. Unlike linear genome sequence, pangenome graphs, and here variation graphs in particular, have the capability to capture the full range of genetic variability characteristic of an evolutionarily or environmentally related collection of genomes (Paten *et al*., 2017). PanTax exploits this quality of variation graphs for capturing the genetic variation that distinguishes the different strains of a species in particular. In addition to this principled quality, pangenome graphs simply offer a considerably more compact way of representing multiple genomes in an encompassing context (Garrison *et al*., 2018; Sirén *et al*., 2021). The practical advantages of pangenome graphs are the preservation of the strain-level diversity of metagenomes and the lack of ambiguity inducing redundancies by which read binning based approaches are affected.

In the following, we first present a comprehensive summary of the workflow of PanTax. Subsequently, we assess its performance at the level of strains in comparison with all prominent state-of-the-art approaches, by experiments using both simulated and real data.

### Approach

Fig. 1 illustrates the workflow of PanTax. Here, we provide an overview of the workflow. For full details, see the Methods section.

**Figure 1.**
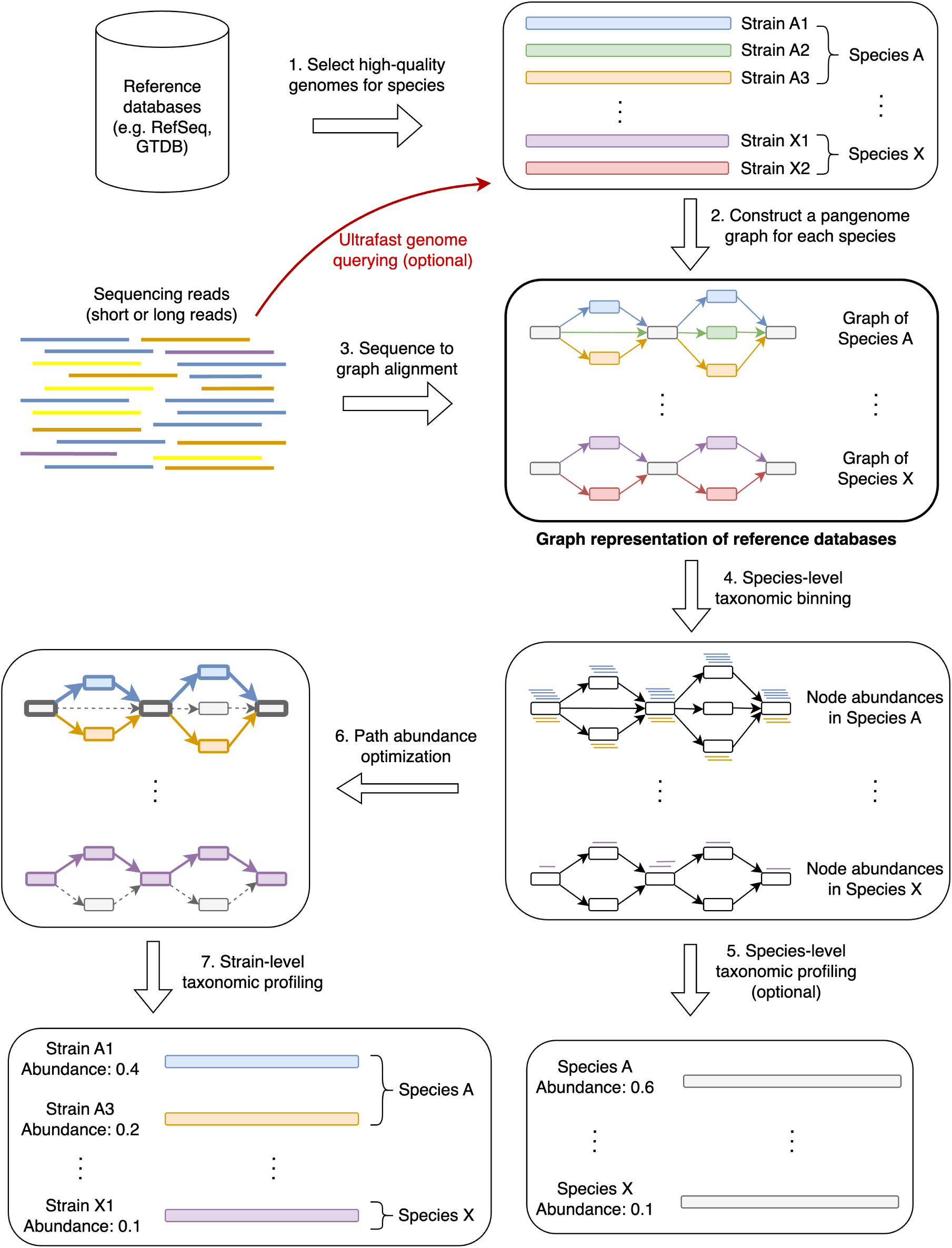
Workflow of PanTax. Different colors indicate different strains. The grey and colored rectangles in the pangenome graphs indicate the shared and unique genomic segments among strains, respectively. Arrows with the same color spell the path of a strain. The bold paths and dashed paths in the pangenome graphs indicate the existent and nonexistent strains in a sample, respectively. For simplicity, only two species (A and X) are shown in the figure. Note that step 5 is optional unless one requires to output the species-level taxonomic profile. The “ultrafast genome querying” step marked with the red arrow and red text indicate the opti^1^o^5^nal way to proceed when running the accelerated version of PanTax (= PanTax (fast)).

PanTax consists of three stages. The first stage is to construct a pangenome graph-based reference database (steps 1-2 in Fig. 1), where each species that makes part of the metagenome corresponds to a species-specific pangenome graph. The second stage (steps 3-5 in Fig. 1) yields species-level taxonomic classification results, by evaluating the amount of reads that gets aligned to each of the species graphs. The third stage eventually determines the strains and the corresponding relative abundances in a species (steps 6-7 in Fig. 1), based on the fact that the individual strains of a species correspond to the individual paths in a species pangenome graph.

From a technical point of view, the first stage reflects to construct a pangenome graph from a collection of high-quality strain-level genomes as stored in reference genome databases such as NCBI’s RefSeq (O’Leary *et al*., 2016) and GTDB (Parks *et al*., 2022), or, if available, in more specific, customized databases. The choice of genome collection can be determined by the user. The more refined collections there are available, the more accurate the results. Note that the flexibility in terms of database usage accounts for the fact that microbe genome databases are filling up rapidly; so, PanTax does not depend on a static snapshot of available microbial sequences, but is explicitly tailored to keep up with the rapid progress in this area of research.

Subsequently, for each read, one needs to determine the species-level graph that gives rise to the optimal read-to-graph alignment of that read. To considerably enhance this step, all species-level pangenome graphs are virtually merged into a large “pangenome super-graph” that captures all species that one has recorded in the first stage. The advantage of merging individual species graphs into one large graph is the fact that one has to align each read with this one large graph, instead of having to align it with each species graph. Apart from drastically decreasing the number of alignment operations, this also facilitates the evaluation of the possibly several alignments of an individual read with different species in a statistically unifying context.

PanTax supports alignment operations for both long and short reads. Based on the resulting alignments, the reads are classified at the level of species. Subsequently, the relative abundance of each of the species is estimated by evaluating the resulting (appropriately normalized) read coverages. Eventually, strain-level profiling can be performed by solving the path abundance optimization problem in each species-specific graph using linear programming (see Methods for definitions). Upon having estimated the abundance of each strain, potentially false positive strains are discarded based on the coverage information of nodes in graphs.

### Datasets

#### Datasets for multi species strain-level taxonomic profiling evaluation tasks

**Simulated datasets (sim-low, sim-high).** We made use of CAMISIM (Fritz *et al*., 2019) to generate 10 metagenomic datasets (2 × 5) with different complexities (i.e. low and high) and different sequencing read types (i.e. Illumina, PacBio HiFi/CLR and ONT R10.4/R9.4.1 reads), which reflect most of application scenarios in current metagenomic research. Here, we applied PBSIM2 (Ono *et al*., 2021), as a most recent tool to simulate PacBio HiFi and ONT reads instead of the default simulator (PBSIM) in CAMISIM using sampling-based simulation mode. The PacBio HiFi sample FASTQ file was down-sampled from an anaerobic digester sludge sequencing sample (SRA accession: ERR7015089) (Feng *et al*., 2022). The ONT R10.4 and R9.4.1 template files are derived from a recent ONT long-read metagenomic study (Sereika *et al*., 2022). The low complexity dataset (named “sim-low”) comprises 60 strains stemming from 30 species (so 2 strains per species on average), while the high complexity (named “sim-high”) dataset comprises 1000 strains from 373 species (so a little less than 3 strains per species on average). For all datasets, the genomes used were retrieved from RefSeq (O’Leary *et al*., 2016)(see Supplementary Data 1 for the details). For the sim-low dataset, the relative abundances of strains vary from 0.30% to 6.43% and corresponding sequencing coverage over different strains varies from 0.96x to 20.3x. The mean sequencing coverage of strains is about 5.3x. For the sim-high dataset, relative abundances of strains vary from 0.04% to 0.30%. The corresponding sequencing coverage over different strains vary from 1.0x to 8.8x, and the mean sequencing coverage of strains is about 2.9x. Notably, in real-world scenarios, some novel strains may be present in a sample, meaning that not all strains can be identified using the reference database. To simulate this situation, we have deliberately included some strains in our simulated samples that are not present in our pangenome reference databases. Specifically, for the sim-low dataset, 56 out of 60 strains are found in the pangenome reference databases, while the remaining 4 strains are not. Similarly, for the sim-high dataset, 795 out of 1000 strains are present in the pangenome reference databases, whereas the other 205 strains are absent.

**Simulated datasets with introduced mutations (sim-low-mut1, sim-high-mut1, sim-low-mut2, sim-high-mut2).** In general, most microbial communities do not contain strains that exactly match reference genomes, as the dominant strains within these communities are often unique to each specific community. Therefore, to reflect this variability, we generated datasets in which the genomes differ from the reference genomes. Building on the genomes used in the previously simulated datasets, we introduced random mutations (including indels and single nucleotide substitutions) into all strains at mutation rates of 0.1% and 1%, thereby creating mutant strains. Using the same read simulation approach, new datasets were generated. Specifically, mutations at 0.1% and 1% were introduced into strains from the sim-low dataset to produce the sim-low-mut1 and sim-low-mut2 datasets, respectively. Similarly, the sim-high dataset was used to generate sim-high-mut1 and sim-high-mut2. In total, considering five types of sequencing data, we generated 20 new metagenomic datasets (2 × 2 × 5) with random mutations.

**Simulated datasets for GTDB (sim-high-gtdb)**. To assess the strain profiling capabilities of PanTax using a large-scale reference database, we generated a high-complexity simulated dataset (sim-high-gtdb) using CAMISIM. This dataset incorporates multiple sequencing technologies, including Illumina, PacBio HiFi/CLR, and ONT R10.4/R9.4.1. Unlike previous simulated datasets that modeled complex microbial communities, the sim-high-gtdb dataset was specifically designed to evaluate PanTax’s performance with a comprehensive reference database. Strain genomes were exclusively sourced from the GTDB:206273 database. Species containing more than two strains were considered, from which 1,000 strains were randomly selected. The sequencing coverage depth of these strains ranged from 1.1× to 9.0×.

#### Real datasets (Zymo1, PD human gut, Omnivorous human gut, Healthy human gut)

*Zymo1.* The ZymoBIOMICS Microbial Community Standard dataset (catalog number: D6330), referred to as Zymo1, comprises sequencing data for Oxford Nanopore Technologies (ONT) R10 reads (https://github.com/LomanLab/mockcommunity, 24 gigabytes FASTQ), ONT R9.4.1 reads (SRA accession: ERR3152364, 27 gigabytes FASTQ), and Illumina reads (SRA accession: ERR2984773, 6.2 gigabytes FASTQ) from an even mock community (Nicholls *et al*., 2019). Note that we used both ONT R10 and R9.4.1 data from Oxford Nanopore GridION sequencing platform. This mock community comprises 10 microbial species: 8 bacteria and 2 yeasts. The relative abundance of each bacteria is 12%, while the relative abundance of each yeast is 2%. As we mainly focus on bacteria in this study, we omitted the two yeasts in our analysis. Therefore, the actual relative abundance of each of the 8 bacteria is re-normalized and designated as 12.5%.

*PD human gut (NGS).* The NGS sequencing data for individuals with Parkinson’s disease (PD) (SRA accession: SRR1906874) were obtained from the study by (Wallen *et al*., 2022). In this investigation, DNA was extracted from fecal samples provided by participants and sequenced using the Illumina NovaSeq 6000 platform. Quality control (QC) of the sequencing data was performed using BBDuk and BBSplit, included in the BBMap package (https://sourceforge.net/projects/bbmap/).

*Omnivorous human gut (PacBio HiFi).* The PacBio HiFi metagenomic sequencing data (SRA accession: SRR17687125, 29.6 gigabytes in FASTQ format) from the fecal sample of a healthy Korean omnivorous individual were sourced from the study by (Kim *et al*., 2022), sequenced on the PacBio Sequel II platform. QC was conducted using fastp (version 0.24.0) (Chen, 2023) and minimap2 (version 2.28-r1209) (Li, 2018). To optimize computational efficiency, 20% of the data were randomly subsampled for analysis using seqtk (version 1.3-r106) (Li *et al*., 2013).

*Healthy human gut (ONT).* The ONT sequencing data (SRA accession: SRR18490940) from the fecal sample of a healthy individual, free of any observable disease or infection, were obtained from the study by (Chen *et al*., 2022). As per the study’s methodology, libraries were prepared using the ONT Ligation library preparation kit (SQK-LSK109, EXP-NBD104, and EXP-NBD114). Sequencing was carried out on the ONT PromethION platform utilizing FLO-PRO002 flow cells. QC was performed using NanoFilt (version 2.8.0) (De Coster *et al*., 2018) and minimap2.

**Spiked-in datasets.** Since accurately evaluating the performance of various methods on the real datasets mentioned above is challenging due to the absence of a known ground truth, we further generated spiked-in datasets to assess these methods in the context of real scenarios. Eight *Vibrio cholerae* strains were selected from a collection of 13,404 complete genomes (as described in Step 1 of the Methods section), and five metagenomic datasets with different sequencing read types (i.e., Illumina, PacBio HiFi/CLR, and ONT R10.4/R9.4.1) were generated using CAMISIM. The ANI between each pair of *Vibrio cholerae* strains ranges from 95.98% to 99.48%. The sequencing coverage of these eight strains varies from 2.4x to 6.0x. The sequencing reads of this eight-strain mixture were then introduced into the corresponding real metagenomic data obtained through small-scale sampling for each sequencing read type. The SRA accessions for the real metagenomic data are as follows: Illumina data (SRR10917775) (Jin *et al*., 2022), PacBio HiFi data (SRR15275213) (Kim *et al*., 2022), PacBio CLR data (SRR10917793) (Jin *et al*., 2022), ONT R10.4 data (SRR27910580) (Campos-Madueno *et al*., 2024), and ONT R9.4 data (SRR18490952) (Chen *et al*., 2022).

#### Dataset for single species strain-level taxonomic profiling evaluation tasks

**Simulated datasets: S. epidermidis strain mixtures (3 strains, 5 strains, 10 strains).** From a total of 34,697 complete genomes in RefSeq (as detailed in Step 1 of the Methods section), we randomly selected 3, 5, and 10 strains of *Staphylococcus epidermidis* (species taxid: 1282) to simulate three NGS metagenomic datasets (corresponding to 3, 5, and 10 strains) using CAMISIM. The ANI between each pair of strains across all three datasets ranges from 99.20% to 99.33%, 98.62% to 99.33%, and 96.96% to 99.47%, respectively. The sequencing coverage of the strains across all three datasets varies from 1x to 10x.

**Real datasets: two cultured S. epidermidis strain mixtures.** In the study by (Emiola *et al*., 2020), the authors cultured two *Staphylococcus epidermidis* strains (species taxid: 1282), NIHLM001 (RefSeq assembly accession: GCF 000276145.1) and NIHLM023 (RefSeq assembly accession: GCF 000276305.1), both isolated from human skin and sharing a 97% average nucleotide identity (ANI). NIHLM001 displayed high resistance to the bacteriostatic antibiotic erythromycin, while NIHLM023 was susceptible. The two strains were mixed in equal proportions (1:1) and cultured under two conditions: one with erythromycin treatment (Ery) and the other without antibiotics (no ATB). Samples were collected at three time points for metagenomic sequencing. For our analysis, we downloaded the sequencing data from the third biological replicate of each condition, resulting in a total of six sequencing datasets.

#### Remarks on evaluating alternative methods

To ensure a rigorous comparison, we have comprehensively selected a variety of representative metagenomic profilers (see Table 1) and evaluated them concurrently. A selection of tools was chosen for evaluation based on the following criteria: open-source availability, active maintenance and/or highly used, acceptable execution time, the ability to create custom databases containing nucleotide sequences, the capability to perform strain-level binning and/or profiling, and good performance in strain classification tasks. Given the principled differences between NGS and TGS reads, we conducted comparisons separately for these two types of reads. For strain-level profiling of multiple species, we ran representative tools such as Centrifuge, Centrifuger, Kraken2, Bracken, Ganon and KMCP on NGS sequencing datasets, while we ran Centrifuge, Centrifuger, Kraken2, Ganon, KMCP and MetaMaps on the TGS sequencing datasets. Note that we excluded CAMMiQ from our benchmarking evaluations because runs ended in segmentation faults in all our attempts.

We ran PanTax in two modes: the default mode and the fast mode. The default mode uses a pre-built pangenome, providing a comprehensive strain profiling. However, the pre-built pangenome requires substantial resources in terms of runtime and memory usage. In contrast, the fast mode first uses the species-level metagenomic profiler (i.e. Sylph) to filter genomes, and then constructs a pangenome tailored to the given samples. This mode greatly reduces resource consumption while still yielding relatively accurate strain profiling results. PanTax operates in two modes: the default mode and the fast mode. The default mode utilizes a pre-built pangenome, offering comprehensive strain profiling. However, this approach demands significant resources in terms of runtime and memory usage. In contrast, the fast mode initially employs efficient strategies (such as those used in Sylph) to filter out potentially false-positive species/strains, followed by the construction of a pangenome incorporating only the remaining samples. This mode significantly reduces resource consumption while still providing strain profiling results that closely approaches those of the default mode.

As for strain-level classification of single species, we compared with the state-of-the-art methods StrainScan, StrainGE and StrainEst, all of which are specifically designed for this task on NGS metagenomic data.

Note that it was observed that usage of default reference databases can introduce systematic biases when using taxonomic classifiers (Simon *et al*., 2019). To mitigate this effect, we utilized a unified set of reference genomes for all methods involved. For strain-level profiling in the multiple species experiments, all methods refer to either RefSeq:13404 or GTDB:206273, because one needs to include all strains into the reference databases for strain-level profiling. RefSeq:13404 encompasses 13,404 strains(8778 species), whereas GTDB:206273 contains 206273 strains (107205 species). We use RefSeq:13404 or GTDB:206273 for constructing pangenomes, because they list several strains for many species and represent commonly used taxonomic classification databases.

### Performance evaluation

To evaluate taxonomic classifiers, we rely on metrics like precision, recall and F1 score, which assess the performance in terms of the identification of taxa (i.e. strains in our case). It is important to realize that these metrics that these metrics are dominated by performance rates on high-abundance taxa. For the sake of an evaluation that does not neglect performance rates on low-abundance taxa, we also utilize precision-recall curves and the area under the curve (AUPR), which capture the desired effects. In addition, accurate estimation of the abundances of the taxa in metagenomes (i.e. profiling) is essential. We use L1 distance, L2 distance and Bray-Curtis (BC) distance to quantify the similarity of the estimated with the true abundance profiles. However, just as before when evaluating performance in binning, these popular distance metrics do not put performance on high-abundance into correct relative context with performance on low-abundance taxa, so tend to neglect the performance on low-abundance taxa. To appropriately account for performance on low-abundance taxa, we also introduce the the complementary metrics absolute frequency error (AFE) and relative frequency error (RFE) so as to capture abundance estimation performance truly comprehensively. See the subsection “Metrics for evaluation” in Methods for full details.

### Tasks

In this study, we focus on two computational tasks, namely, strain-level taxonomic profiling for multiple species and for single species. Strain-level taxonomic profiling for multiple species provides a detailed view of strain diversity and abundance across different species, enhancing our understanding of microbial dynamics. Additionally, strain-level profiling for a single species allows for in-depth analysis of strain variation within that species. We evaluate PanTax and other state-of-the-art methods using both NGS and TGS data from various datasets to ensure comprehensive benchmarking. See below for the details.

### Strain level taxonomic profiling for multiple species

**Simulated datasets.** Figure 2 presents the benchmarking results. For the sake of a clear presentation, Figure 2 only focuses on the most important metrics. For detailed results, referring to all metrics, please see Supplementary Tables 1,2, corresponding to the two datasets sim-low and sim-high. In terms of taxon identification, when compared with all other tools, PanTax achieves high precision across all sequencing data types, including NGS, PacBio HiFi/CLR, and ONT R9.4.1/R10.4. Specifically, for NGS, PacBio CLR, and ONT R10.4 data, PanTax achieves the highest precision (sim-low/sim-high: 0.918/0.806, 0.889/0.778, 0.918/0.826) across both sim-low and sim-high datasets. On both sim-low and sim-high datasets (PacBio HiFi, ONT R9.4.1), KMCP achieves the highest precision (0.960/0.884, 1.000/0.950) at the cost of very low recall (0.400/0.479, 0.317/0.362), while PanTax achieves the second highest precision (0.948/0.854, 0.889/0.779) while maintaining comparable recall with other methods. Overall, PanTax achieves the highest F1 score for both sim-low and sim-high datasets across all sequencing data types, while maintaining comparable AUPR compared to other tools. Notably, PanTax (fast) even outperforms PanTax in terms of precision, particularly on sim-high datasets. However, for datasets with high sequencing error rates, such as PacBio CLR and ONT R9.4.1, the recall of PanTax (fast) tends to decrease. This is likely due to the “ultrafast genome querying” step, which fails to detect the presence of some strains as noisy reads may bias k-mer containment.

**Figure 2.**
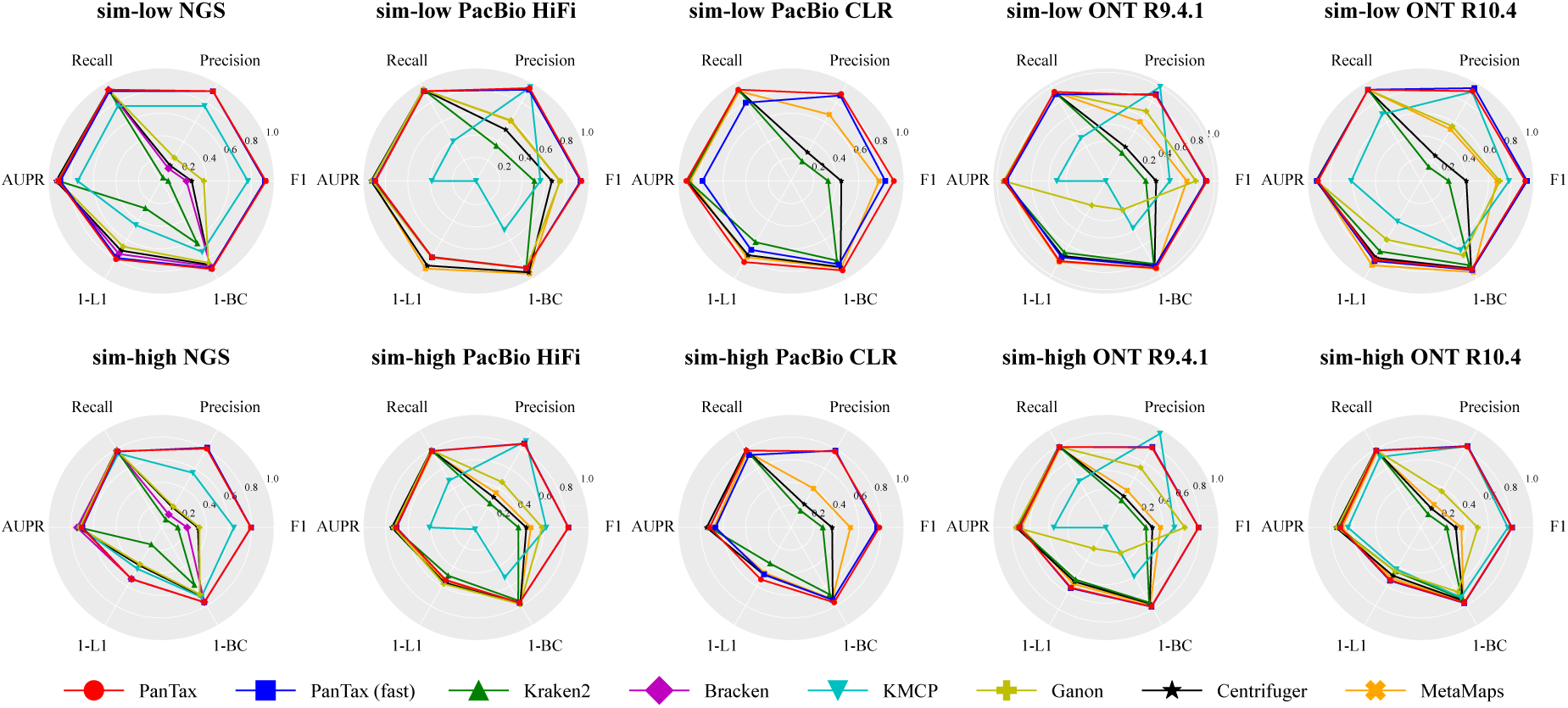
Benchmarking results of strain-level taxonomic profiling on the simulated datasets (sim-low and sim-high). The upper and lower panels display the sim-low and sim-high datasets, respectively, for each of the five sequencing read types. AUPR: area under the precision-recall curve. To visualize all metrics consistently (i.e., with higher values indicating better performance), we present the 1-L1 distance and 1-BC distance.

For evaluating taxonomic abundances, PanTax (0.100/0.237) demonstrates superior performance in terms of BC distance for NGS datasets, outperforming other methods across both simulated datasets. On sim-low datasets (PacBio HiFi, ONT R9.4.1/R10.4), MetaMaps achieves the best BC distance, but it performs worse than PanTax on PacBio CLR data. PanTax achieves the second best BC distance on the ONT R9.4.1/R10.4 sim-low datasets. Conversely, on sim-high datasets (PacBio CLR, ONT R9.4.1/R10.4), PanTax achieves the best BC distance performance compared with other methods. Ganon performs the best for BC distance on the sim-high dataset (PacBio HiFi). Notably, PanTax (fast) achieves comparable BC distance to PanTax, except on error-prone datasets such as PacBio CLR and ONT R9.4.1 data. Additionally, similar performance trends are observed for PanTax on other related metrics such as L1 and L2 distances, AFE, and RFE.

**Simulated datasets with introduced mutations.** Figure 3 displays the benchmarking outcomes for strain-level taxonomic profiling across multiple simulated datasets with introduced mutations. Detailed results for all metrics can be found in Supplementary Tables 3,4,5,6, corresponding to the four datasets sim-low-mut1, sim-high-mut1, sim-low-mut2, sim-high-mut2.

**Figure 3.**
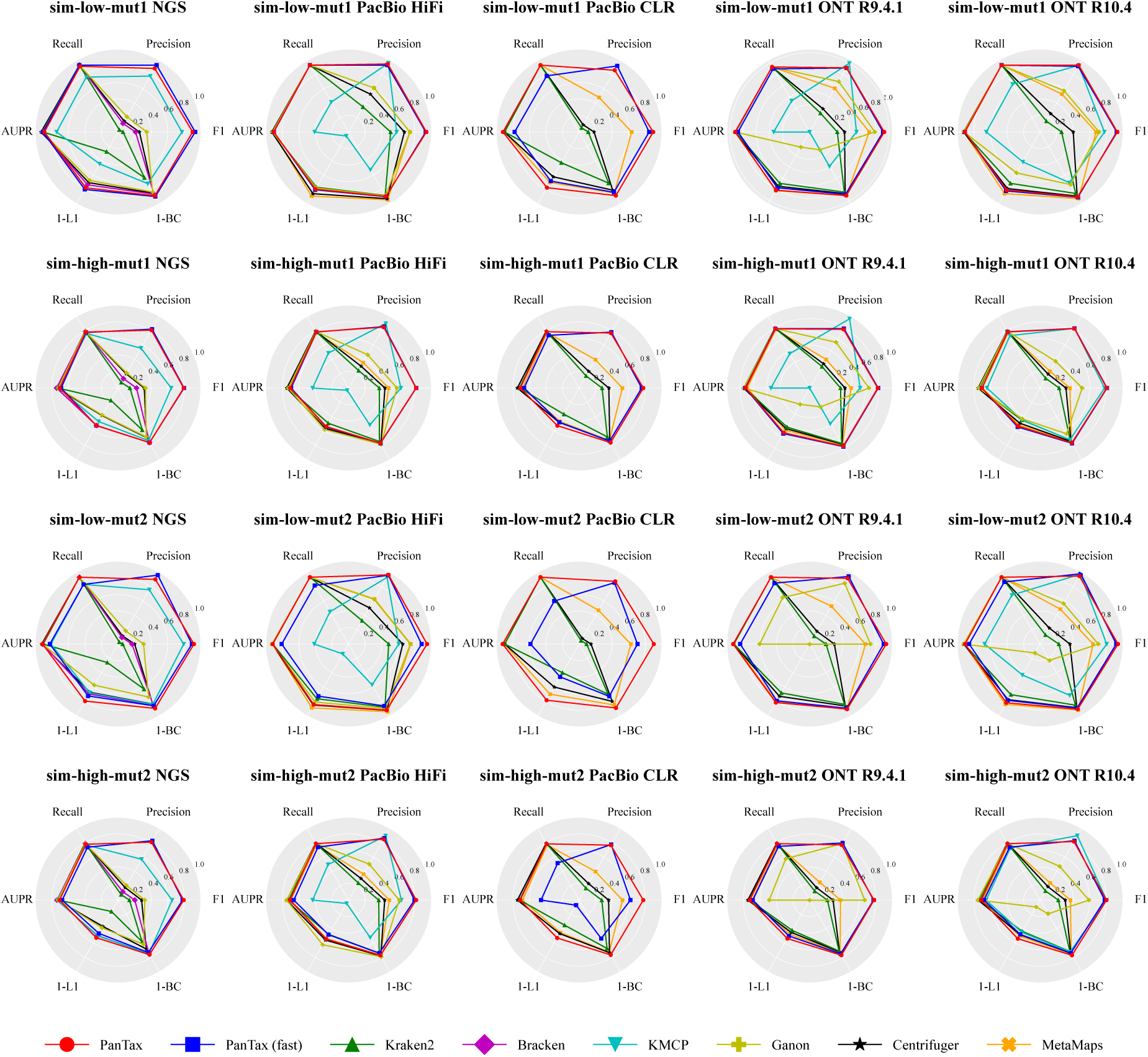
Benchmarking results of strain-level taxonomic profiling on the simulated datasets with introduced mutations (sim-low-mut1, sim-high-mut1, sim-low-mut2, sim-high-mut2). AUPR: area under the precision-recall curve. To visualize all metrics consistently (i.e., with higher values indicating better performance), we present the 1-L1 distance and 1-BC distance.

For datasets with 0.1% mutations (sim-low-mut1 and sim-high-mut1), PanTax demonstrates notably high precision compared to other tools across all sequencing types, including NGS, PacBio HiFi/CLR, and ONT R9.4.1/R10.4. Specifically, for NGS and PacBio CLR, PanTax achieves the highest precision (0.887/0.808, 0.862/0.768) across both simulated datasets, while for ONT R10.4 datasets, PanTax achieves the highest and the second highest precision (0.933/0.829) in both simulated datasets, respectively. In both sim-low-mut1 and sim-high-mut1 datasets (PacBio HiFi, ONT R9.4.1), KMCP reaches the highest precision (0.962/0.902, 1.000/0.957), but with a significant trade-off in recall, which is substantially lower. PanTax, on the other hand, achieves the second-highest precision (0.949/0.850, 0.918/0.775) while maintaining comparable recall to other methods. In terms of overall performance, PanTax achieves the highest F1 score across all sequencing types for both sim-low and sim-high datasets, while maintaining similar AUPR to other tools. Regarding taxonomic abundance estimation, for NGS datasets, PanTax (0.110/0.238) performs better in terms of BC distance than the other methods across both simulated datasets. On sim-low-mut1 datasets (PacBio HiFi, ONT R9.4.1/R10.4), MetaMaps performs best in terms of BC distance but underperforms compared to PanTax on PacBio CLR data. On sim-high-mut1 datasets (PacBio CLR, ONT R9.4.1/R10.4), however, PanTax achieves the best BC distance performance compared to the other methods.

For datasets with 1% mutations (sim-low-mut2 and sim-high-mut2), PanTax maintains the highest precision across all sequencing types for sim-low-mut2 datasets and outperforms other tools in terms of precision for NGS, PacBio CLR, and ONT R10.4 sequencing types on sim-high-mut2 datasets. In addition, PanTax achieves the highest F1 score for both sim-low-mut2 and sim-high-mut2 datasets across all sequencing data types. In evaluating taxonomic abundances, for NGS, PacBio CLR, and ONT R9.4.1 datasets, PanTax (0.102/0.238, 0.108/0.235, 0.092/0.231) outperforms the other methods in terms of BC distance across both simulated datasets. On sim-low-mut2 datasets (PacBio HiFi, ONT R10.4), MetaMaps achieves the best BC distance. However, on sim-high-mut2 datasets (NGS, PacBio CLR, ONT R9.4.1/R10.4), PanTax achieves the best BC distance performance. Ganon achieves the best BC distance on the sim-high-mut2 dataset (PacBio HiFi), but with only a slight advantage over PanTax. It is important to note that the comparison between PanTax (fast) and PanTax reveals similar patterns across most metrics, as observed in the original sim-low and sim-high datasets.

**Real datasets.** To assess performance on the Zymo1 mock community datasets, we specifically constructed a new reference database to encompass most of the strains in Zymo1. This reference database includes only eight species, but each species consists of a greater number of strains selected from 34,697 complete genomes in RefSeq (as described in Step 1 of the Methods section). The number of strains per species is limited to 80 due to aligner performance constraints. All benchmarking tools use this reference database to ensure a fair comparison. As shown in Figure 4, the strain-level taxonomic profiling results reveal that PanTax (0.875/0.438/0.500) significantly outperforms other methods in terms of precision on NGS, ONT R9.4.1, and R10.4 data. Furthermore, PanTax achieves the highest F1 score across NGS, ONT R9.4.1, and R10.4 data. Bracken achieves the best AUPR on NGS data, while Kraken2 obtains the best AUPR (0.858) on ONT R9.4.1 data, and MetaMaps achieves the best AUPR (0.751) on ONT R10.4 data. Nonetheless, PanTax also achieves the second-best AUPR (0.816/0.748) on ONT R9.4.1/R10.4 data. Regarding the evaluation of taxonomic abundances, PanTax shows the lowest BC distance on NGS data, while MetaMaps achieves the lowest BC distance on ONT data. Additionally, PanTax demonstrates the lowest L2 distance across various data types, including NGS and ONT. PanTax significantly outperforms other methods in terms of AFE and RFE across both NGS and ONT sequencing data. Note that detailed results for all metrics can be found in Supplementary Table 7. To evaluate the feasibility of PanTax on real metagenomic data, we assessed its performance alongside several leading metagenomic profilers (according to BC distance in our simulated and mock community datasets) using human gut microbiome datasets, including PD human gut (NGS), Omnivorous human gut (HiFi), and Healthy human gut (ONT) datasets. PanTax was compared with Centrifuger, Kraken2, Bracken, and KMCP on the NGS dataset, while comparisons on the HiFi and ONT datasets included Centrifuger, Kraken2, Ganon, and MetaMaps. Given the absence of ground truth for real human gut metagenomes, we focused on comparing the relative taxon abundances of commonly detected strains across PanTax and other profilers. As shown in Figure 5, PanTax demonstrates a strong correlation with other state-of-the-art metagenomic profilers in terms of taxon abundance. Specifically, the Pearson correlation coefficient reaches 0.95 with Bracken on the NGS dataset, 0.86 with MetaMaps on the HiFi dataset, and 0.97 with MetaMaps on the ONT dataset. Notably, Bracken and MetaMaps are among the most competitive methods across most simulated and mock community datasets. In summary, these results highlight the capability of PanTax to perform strain-level taxonomic profiling on real, complex metagenomic samples.

**Figure 4.**
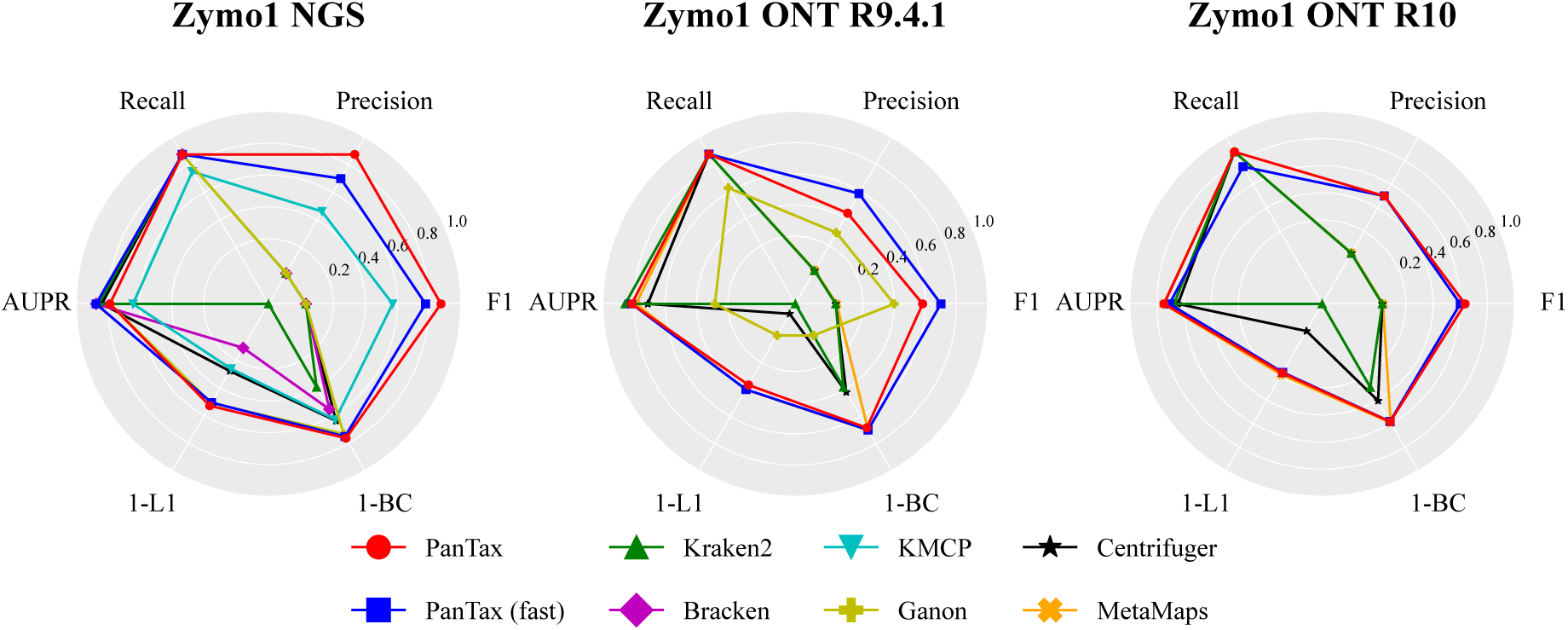
Benchmarking results of strain-level taxonomic profiling on the real datasets (Zymo1 mock community). AUPR: area under the precision-recall curve.To visualize all metrics consistently (i.e., with higher values indicating better performance), we present the 1-L1 distance and 1-BC distance.

**Figure 5.**
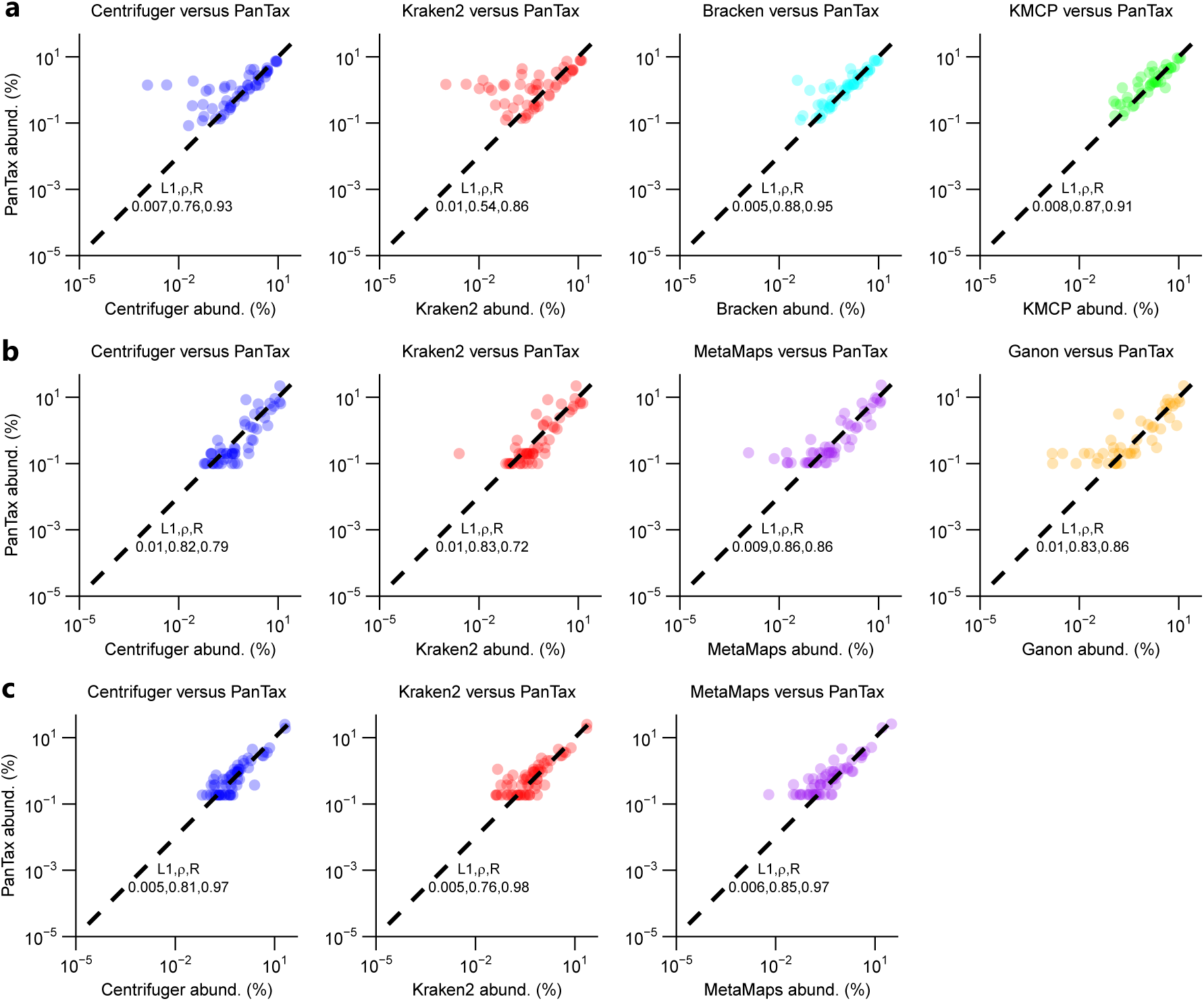
Results of strain-level taxonomic profiling for real human gut metagenomes. **(a)** PD human gut (NGS) dataset, **(b)** Omnivorous human gut (PacBio HiFi) dataset, and **(c)** Healthy human gut (ONT) dataset. For all three datasets, the relative taxon abundance correlation between PanTax and other competitive profilers was computed. The comparisons included mean L1 distance, Spearman correlation, and Pearson correlation. Note that we failed to run Ganon and KMCP on the Healthy human gut (ONT) dataset because it was primarily designed for NGS data.

**Spiked-in datasets.** Figure 6 presents the benchmarking results for strain-level taxonomic profiling on the spiked-in metagenomic datasets. For NGS data, PanTax and KMCP achieve the highest precision, recall, F1 score, and AUPR (all equal to 1.0). Bracken performs best in terms of abundance estimation, as indicated by metrics such as BC/L1/L2 distances, AFE, and RFE, although PanTax shows only slightly worse performance than Bracken and KMCP. For HiFi data, PanTax, Ganon, Centrifuge, and Centrifuger achieve the highest precision, recall, F1 score, and AUPR (all equal to 1.0). Centrifuger excels in abundance estimation, but PanTax still outperforms Ganon in this aspect. Most importantly, for handling noisy reads, such as those from PacBio CLR and ONT R9.4.1/R10.4 data, PanTax substantially outperforms all other tools across all metrics. As these spiked-in datasets reflect both strain diversity (with an 8-strain mixture) and the complexity of real metagenomic samples, the results demonstrate the advantages and robustness of PanTax for metagenomic profiling on complex, real-world samples. Note that detailed results for all metrics can be found in Supplementary Table 8.

**Figure 6.**
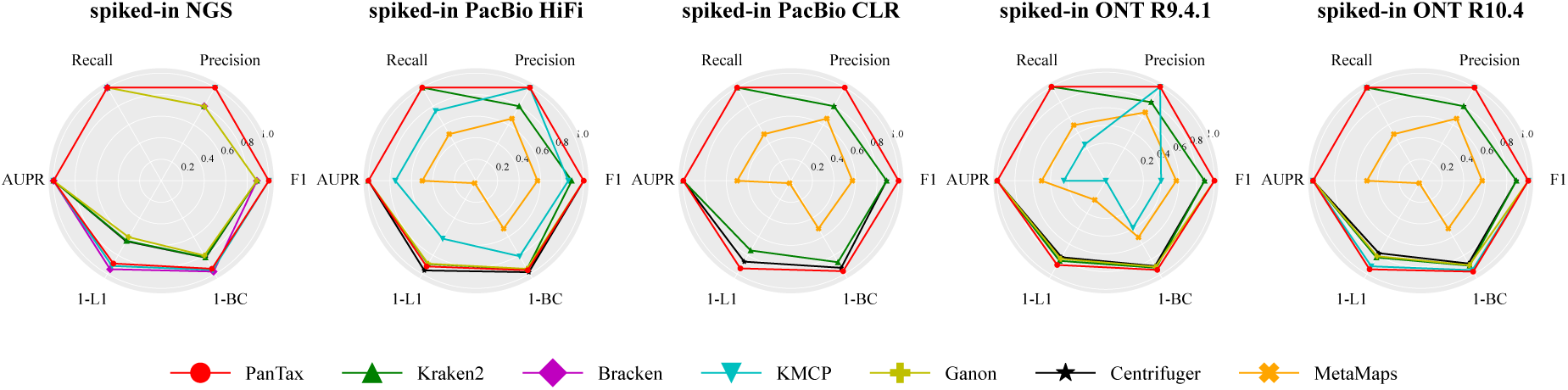
Benchmarking results of strain-level taxonomic profiling on the spiked-in datasets. AUPR: area under the precision-recall curve. To visualize all metrics consistently (i.e., with higher values indicating better performance), we present the 1-L1 distance and 1-BC distance.

### Strain level taxonomic profiling for single species

We note that several prominent metagenomic tools for strain-level taxonomic classification, such as Centrifuger, Bracken, Kraken2, KMCP, Ganon and MetaMaps, have the capability to identify multiple species or strains and determine their relative abundances within a metagenomic sample. In addition to these versatile tools, there are others specifically designed to generate strain-level abundance profiles but are tailored for single-species analysis and NGS data. These tools require users to preassign the species they wish to analyze, after which they provide outputs detailing the abundances of strains within the specified species. Noteworthy examples of such tools include StrainScan, StrainGE, and StrainEst (see Table 1).

To compare PanTax with other methods capable of performing strain-level microbiome composition analysis for single species from NGS reads, we conducted several benchmarking experiments.

**Simulated datasets: S. epidermidis strain mixtures (3 strains, 5 strains, 10 strains).** For the simulated datasets, we selected *Staphylococcus epidermidis* (species taxid: 1282) as an example. It is important to note that when dealing with a single species, PanTax faces challenges in constructing a pangenome from an excessive number of individual genomes (e.g., *>* 90). Therefore, we applied clustering techniques to remove highly similar (redundant) genomes before constructing the pangenome. Other methods also incorporate built-in techniques for eliminating redundant genomes, ensuring the fairness of our comparison.

First, we chose all complete genomes (147) of *S. epidermidis* from the RefSeq database and subsequently obtained 70 non-redundant genomes (strains) of this species using single-linkage clustering with a threshold of ANI ≤ 99.9%. These genomes were subsequently utilized to construct the reference pangenome graph for this species. As shown in Figure 7 (detailed values of each metric are provided in

**Figure 7.**
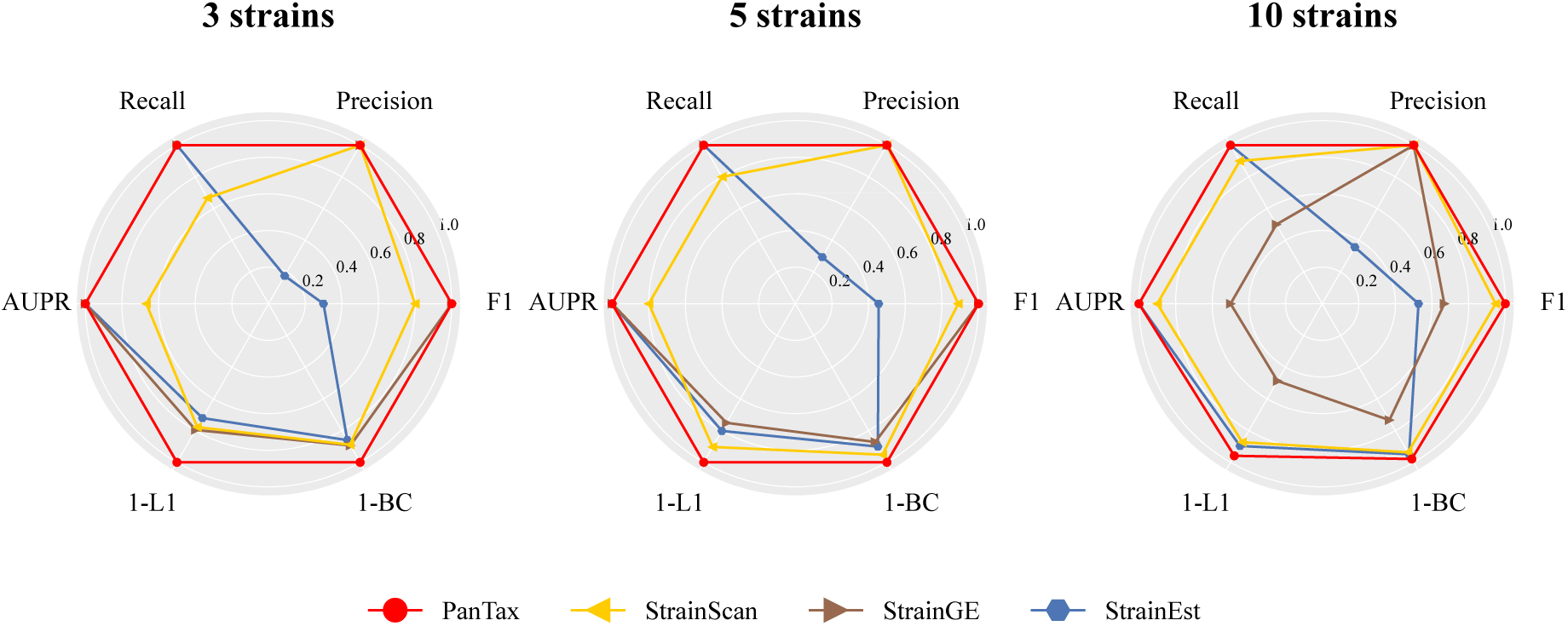
Benchmarking results of single species strain-level taxonomic profiling on the simulated datasets (*S. epidermidis* strain mixtures:3 strains, 5 strains, 10 strains). AUPR: area under the precision-recall curve. To visualize all metrics consistently (i.e., with higher values indicating better performance), we present the 1-L1 distance and 1-BC distance.

Supplementary Table 9), PanTax achieved nearly perfect performance on all three datasets, attaining 100% precision, recall, and AUPR, as well as nearly zero BC distance. This significantly outperforms other state-of-the-art approaches, such as StrainScan, StrainGE and StrainEst.

**Real datasets: antibiotic-resistant S. epidermidis strain mixtures (2 strains).** Similarly, we selected a total of 1395 *S. epidermidis* genomes (including both complete and incomplete genomes) from the RefSeq database, as these two strains are not included in the complete reference genomes.

To handle more reference genomes in this case, we employed a more effective strategy for removing redundant genomes. This strategy consists of two stages. In the first stage, we performed complete-linkage clustering using an ANI threshold of 99% to reduce redundancy, resulting in 69 non-redundant genome clusters. Representative genomes from these clusters were then selected as the reference genomes, marking the completion of the first redundancy removal stage. In the second stage, we ran PanTax with less stringent parameter settings (*f*_strain_ set to 0.2) to filter out clusters where the representative genome was not identified by PanTax. We then merged all genomes from the remaining clusters and applied a more stringent ANI threshold of 99.9% for graph-based clustering approach (see Algorithm 1 in the Supplementary Methods) to further reduce redundancy, similar to the approach used for the previous simulated datasets. The difference is that all non-redundant genomes were selected, rather than the default number of 10 non-redundant genomes. After completing these two stages of redundancy removal, we selected the non-redundant genomes as the final reference genomes to construct the pangenome graph, which was used as input for running PanTax with more stringent parameter settings (*d*_strain_ set to 0.25). We compared PanTax with other tools that perform strain-level taxonomic classification for these *S. epidermidis* 2-strain mixtures, with the results shown in Figure 8. These mixtures consist of two *S. epidermidis* strains, among which GCF 000276305.1 is not resistant to the antibiotic erythromycin, while GCF 000276145.1 exhibits high resistance to erythromycin, and their relative abundances vary at different time points, as reported in the original study (Emiola *et al*., 2020). GCF 000276305.1 was consistently the dominant strain in the no ATB group, while GCF 000276145.1 was the dominant strain in the Ery group at each time point. The results demonstrate that both PanTax and StrainScan correctly identified the two strains in all six samples. Furthermore, PanTax and StrainScan showed similar relative abundance estimations. StrainGE and StrainEst returned two main correct representative strains. However, the relative proportions of the two dominant strains detected by StrainGE in the no ATB group do not match the ground truth. StrainGE identified GCF 012029805 as the dominant strain instead of GCF 000276305 in the no ATB group. StrainEst consistently detected false-positive minor strains.

**Figure 8.**
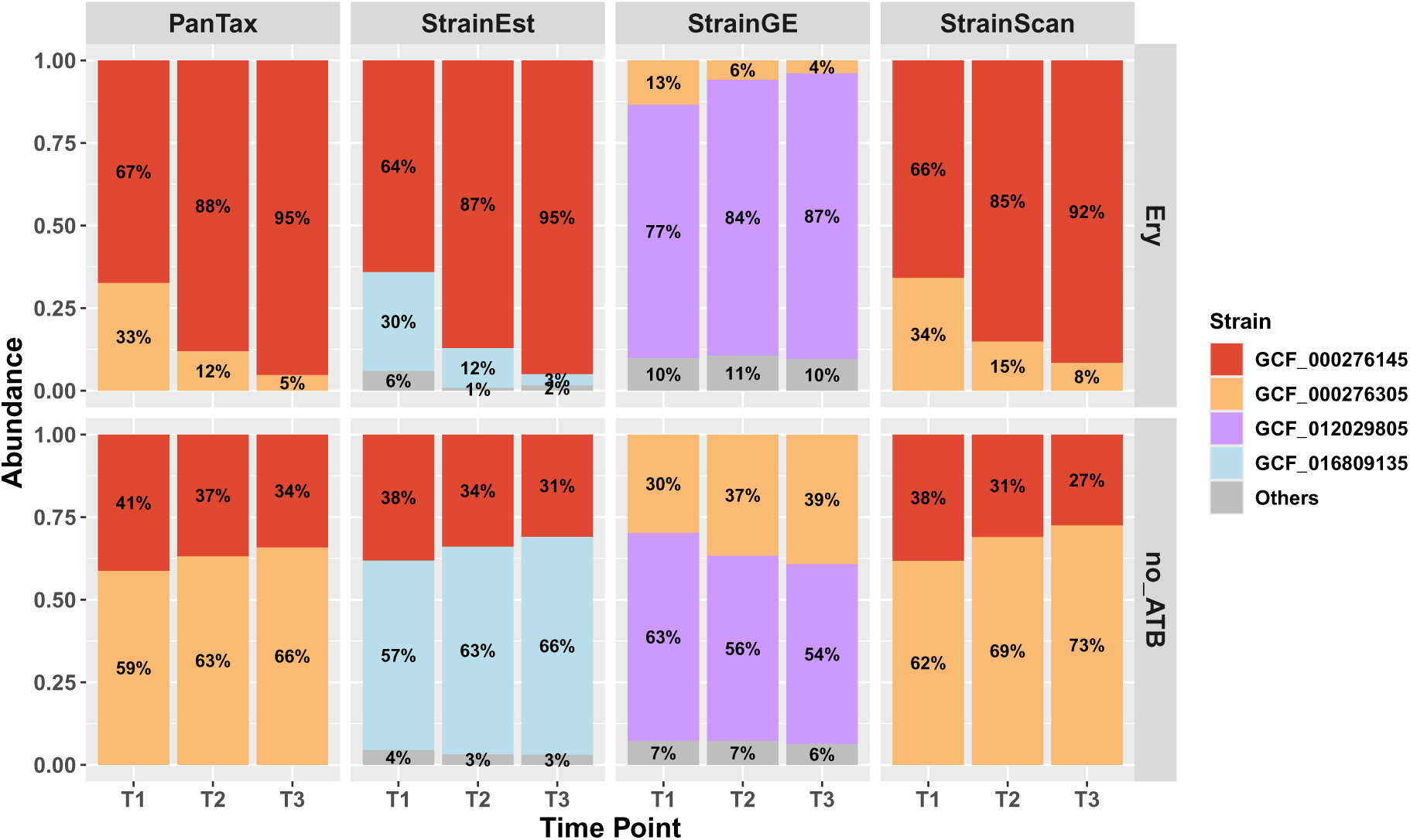
Benchmarking results of single species strain-level taxonomic profiling on real datasets: two-cultured *S. epidermidis* strain mixtures. The x-axis represents time points, and the y-axis shows the relative taxon abundance of each identified strain. The two strains (marked in red and yellow) were mixed in equal proportions (1:1) and cultured under two conditions: one with erythromycin treatment (Ery) and one without antibiotics (no ATB). The upper and lower panels display the strain abundances predicted by four tools under Ery and no ATB, respectively.

### Effects of divergence versus ratio of abundances

To explore the impact of divergence and abundance disparities on strain identification in mixed samples, we simulated multiple mixtures comprising two strains. Similarly, we used the reference database for *Staphylococcus epidermidis* (species taxid: 1282) which was previously constructed for the Staphylococcus epidermidis simulated datasets and selected 6 strains from it to perform the experiment. These mixtures encompassed various combinations of ANI values of 96.8%, 97%, 98%, 99%, and 99.8% (with 99.8% representing the most challenging scenario and 96.8% the least) and abundance ratios of 1:1, 1:3, 1:5, and 1:10. Our experiments centered on NGS reads, and we subsequently assessed performance of PanTax using F1 score, AUPR, and L2 distance for each possible combination of divergence and abundance ratio. The results are presented in Supplementary Figure 1. Analysis of the F1 score and AUPR indicates that PanTax successfully identifies both true strains in all scenarios. Regarding L2 distance, while PanTax accurately estimates the relative abundances of each strain in most cases, it exhibits biased abundance estimations for the most challenging scenario (ANI=99.8%).

### Effects of reducing sequencing coverage

To study the effects of reducing sequencing coverage, we randomly subsampled 1/2 and 1/5 of the reads from the “sim-low” datasets, naming these subsets “sim-low-sub1” and “sim-low-sub2”, respectively. We then benchmarked all related methods accordingly. Supplementary Tables 10 and 11 present the strain-level taxonomic classification results for the sim-low-sub1 and sim-low-sub2 datasets, respectively. For sim-low-sub1, PanTax significantly outperforms other methods (with the exception of KMCP) in strain-level precision across most sequencing data types, including NGS, PacBio HiFi, and ONT R9.4.1/R10.4. However, KMCP exhibits considerably lower recall, resulting in PanTax or PanTax (fast) achieving the highest F1 score across these sequencing technologies. Additionally, PanTax demonstrates comparable or superior performance in taxonomic abundance estimation, as indicated by metrics such as L2 distance, BC distance, AFE, and RFE. Notably, for PacBio CLR data, PanTax attains the second-highest precision, trailing behind MetaMaps. Nevertheless, it achieves superior recall and AUPR while maintaining comparable L2 distance and AFE relative to MetaMaps. A similar trend is observed for sim-low-sub2, where PanTax or PanTax (fast) again outperforms other methods (except KMCP) in strain-level precision across all sequencing data types, including NGS, PacBio HiFi/CLR, and ONT R9.4.1/R10.4. However, KMCP exhibits substantially lower recall, leading to a significantly reduced F1 score, particularly in TGS datasets. Regarding taxonomic abundance estimation, PanTax or PanTax (fast) achieves performance comparable to or slightly below that of the top-performing tools. As sequencing coverage decreases, PanTax exhibits lower recall but higher precision in strain identification. Additionally, its taxonomic abundance estimation becomes less accurate. In summary, these experiments demonstrate that PanTax or PanTax (fast) is the superior choice of approach in particular when dealing with low coverage (components of) metagenomes.

### PanTax is effective for larger reference metagenome databases

To demonstrate that PanTax is capable of handling larger reference metagenome databases, such as the Genome Taxonomy Database (GTDB, Release R220, containing 206,273 strains)(Parks *et al*., 2022), we specifically developed an optimized version, PanTax (fast), as detailed in the Methods section. We benchmarked PanTax (fast) against other approaches using the sim-high-gtdb datasets (as reflecting complex metagenomes) across all five sequencing data types (using GTDB:206273). The results are shown in Figure 9 (detailed values of each metric are provided in Supplementary Table 12). For NGS data, PanTax (fast) achieves the second-best precision (0.995), which is only slightly lower than KMCP (1.0). While PanTax (fast) shows lower recall compared to other methods, its BC distance is significantly better than tools like Ganon, Centrifuge, and Kraken2. For the four TGS datasets, KMCP is the only competitor that outperforms PanTax (fast) in precision, albeit with a very slight advantage(0.001 to 0.005). However, PanTax (fast) substantially surpasses KMCP in other metrics, including recall (0.097 to 0.467 higher), AUPR, L1, and BC distances. In summary, PanTax (fast) demonstrates comparable overall performance to KMCP on NGS data but substantially outperforms all other methods on TGS data.

**Figure 9.**
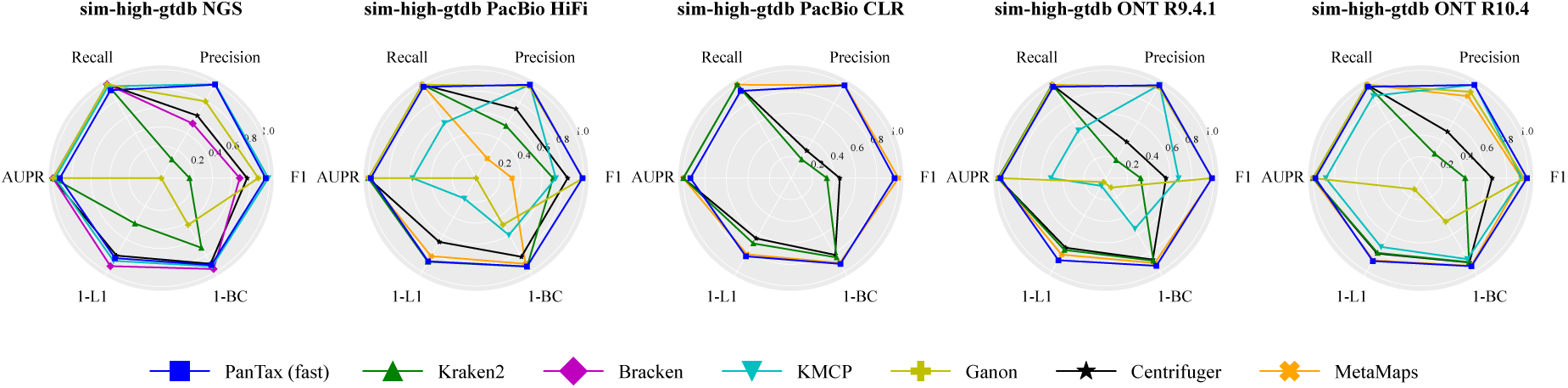
Benchmarking results of strain-level taxonomic profiling on the simulated datasets (sim-high) using GTDB database. Only the results of PanTax (fast) are shown, as the original PanTax is too time-consuming to run using GTDB. AUPR: area under the precision-recall curve. To visualize all metrics consistently (i.e., with higher values indicating better performance), we present the 1-L1 distance and 1-BC distance.

### Runtime and memory usage evaluation

Supplementary Tables 13 and 14 present the runtime and peak memory usage of benchmarking tools using NGS and TGS datasets, respectively, with the RefSeq:13404 reference database. We primarily compare the runtime and memory usage for index construction and read alignment during queries, as these processes dominate the main computational resources. When dealing with NGS reads, PanTax initially requires index construction for the pangenome graphs, whereas this indexing process is unnecessary for TGS reads. Notably, the indexing process accounts for the majority of the time compared to the read-to-graph alignment. However, for a given pangenome reference, indexing needs to be performed only once. Notably, here and in the following sections, the taxonomic profiling time for all tools refers to the total time taken to execute the tools, including the time required for database construction. PanTax exhibits longer database construction and profiling times and requires larger peak memory usage compared to other tools. However, PanTax (fast) achieves comparable performance in terms of time and superior performance in terms of memory usage, when compared to other tools. For example (see Figure 10), on the sim-low dataset, PanTax (fast) requires 10.5 CPU hours, making it faster than tools such as KMCP (19.3 hours), Kraken2 (567 hours), and Centrifuge (152.6 hours), and only slightly slower than Ganon (7.5 hours) and Bracken (7.8 hours). Additionally, PanTax (fast) requires the lowest peak memory usage (17.9 GB), whereas the second-lowest memory tool, KMCP, requires 18.8 GB. In the case of the real PD human gut dataset, PanTax (fast) consumes 29.7 CPU hours, which is only slower than Ganon (7.2 hours) and Bracken (7.8 hours), but several times faster than other tools such as Kraken2, Centrifuge, and Centrifuger. Furthermore, it requires 14.9 GB of RAM, which is the lowest memory usage among all the tools.

**Figure 10.**
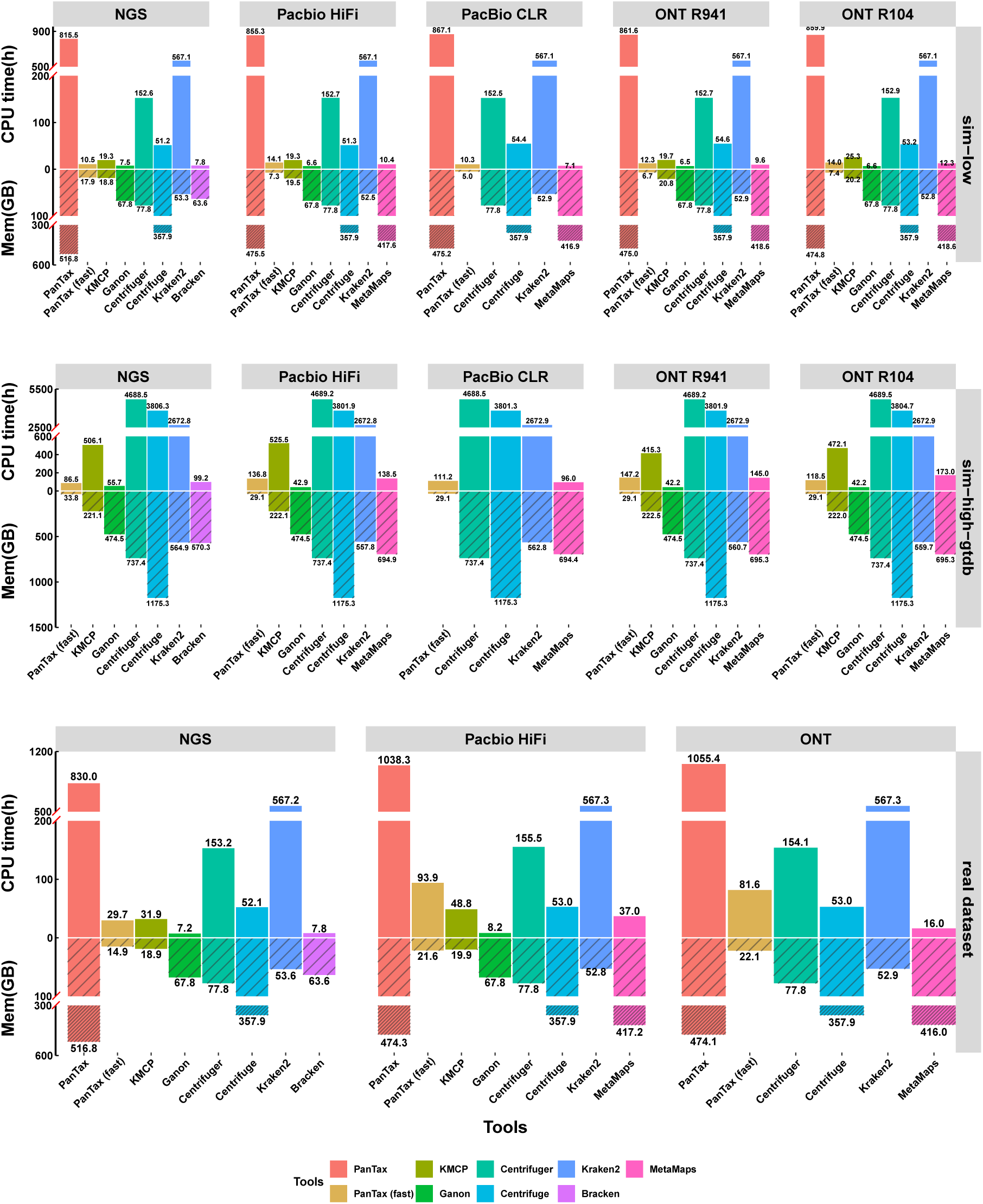
The CPU time and peak memory usage (Mem) of various metagenomic profilers. The upper, middle, and bottom panels represent the sim-low, sim-high-gtdb, and real human gut datasets, respectively. Note that for the real datasets, from left to right, the panels correspond to the PD human gut (NGS), Omnivorous human gut (PacBio HiFi), and Healthy human gut (ONT) datasets.

Similar to NGS datasets, in TGS datasets, PanTax exhibits longer database construction and profiling times and requires higher peak memory usage compared to other tools (see Supplementary Table 14). However, widely used tools such as Kraken2 and MetaMaps require comparable time and memory usage, respectively. Despite this, PanTax (fast) delivers performance that is either superior or comparable to other tools (see Supplementary Table 14). For example (see Figure 10), on sim-low (ONT R9.4.1) data, PanTax (fast) requires 12.3 CPU hours, making it slightly slower than the fastest two tools (Ganon: 6.5 hours, MetaMaps: 9.6 hours), but MetaMaps consumes a substantial amount of memory (around 400 GB) whereas Ganon also requires a moderate amount of memory (37 GB). In contrast, PanTax (fast) requires the lowest memory usage (6.7 GB), with the second lowest being KMCP (20.8 GB). On the real Omnivorous human gut dataset (HiFi), PanTax (fast) requires 93.9 CPU hours, which is slower than Ganon (8.2 hours), KMCP (48.8 hours), Centrifuge (53 hours) and MetaMaps (37 hours), but shows 1.7× and 6× faster than Centrifuger and Kraken2, respectively. Regarding memory usage, PanTax (fast) requires comparable memory usage to KMCP (21.6 GB and 19.9 GB, respectively), both of which require 5 to 20 times less memory than the alternative methods. On the sim-high-gtdb datasets, which utilize a large-scale reference database (GTDB:206273), PanTax (fast) achieves comparable speed to MetaMaps, being only 1-3 times slower than Ganon, but significantly faster (approximately 4-50 times) than other tools such as KMCP, Kraken2, Bracken, Centrifuge, and Centrifuger (see Figure 10 and Supplementary Table 15). Notably, PanTax (fast) requires the lowest peak memory usage (around 30 GB) across all sim-high-gtdb datasets, while other tools require at least 200 to 1200 GB (see Figure 10 and Supplementary Table 14).

It is worth noting that for profiling a single species, although PanTax requires more time and memory than other tools like StrainScan, StrainGE, and StrainEst, the time (approximately 27 CPU hours) and memory (around 20 GB) required are still acceptable (see Supplementary Table 15).

## Discussion

We have introduced PanTax, an approach that classifies the contents of metagenomes in terms of their taxa. PanTax accepts all classes of popular sequencing reads as input, and determines the organisms that make part of the metagenome at the level of their strains. While classification of the contents of metagenomes at the level of species had reached a satisfying level of maturity in the literature, determining presence and abundance of strains of species in metagenomes has still remained in its infancy. Correspondingly, PanTax addresses this important open challenge.

Beyond delivering decisive progress with respect to the characterization of the strain content of metagenomes, PanTax also delivers relevant practical advantages. Namely, PanTax is not prone for biases relative to pre-selected genomic regions, PanTax can process all popular types of reads, it can handle multiple species simultaneously, and it can integrate custom databases. The latter, in particular, is very important, because it ensures to keep up with the already fast space in terms of the detection of novel bacterial species and strains. Last but not least, PanTax also provides sound estimates on the abundances of the different taxa detected.

Only two alternative approaches, Centrifuge and Centrifuger, account for the same level of comprehensiveness when evaluating metagenomes. Experiments on simulated and real data demonstrate that PanTax substantially outperforms these two prior approaches, both of which gather all of the qualities that PanTax boasts, substantially. Beyond outperforming all approaches that account for similar comprehensiveness, PanTax also outperforms or is on a par with the state-of-the-art that specializes in particular, specific aspects of the taxonomic classification of metagenomes.

Key to success is the fact that PanTax, to the best of our knowledge, is the first approach that builds on pangenome graphs as a foundation in terms of reference systems, instead of linear genomes. Unlike ordinary linear reference genomes, pangenome graphs establish reference systems that can capture the diversity inherent to a collection of genomes even at the level of strains, the finest taxonomic resolution possible. The type of pangenome graph that PanTax is based on are variation graphs. Beyond just capturing the diversity of a mix of genomes, variation graphs also arrange all genomes in an evolutionarily consistent manner, which eliminates ambiguities during classification.

This lack of ambiguities is particularly beneficial when determining the strain or species specific origin of single reads: because a read aligns with a graph that incorporates all genomes evolutionarily consistently, the read-to-graph alignment immediately points out the strain (and, of course, also the species) the read stems from. Instead, aligning a read against the applicable linear reference genomes entails a statistically involved analysis of the alignment scores that each of the different alignments delivers; because the single linear genomes do not explicitly relate with each other, this analysis may leave one with statistical uncertainties that are hard to resolve. A particularly interesting scenario results from the failure of a read to align with any of the linear reference genomes. Still, however, the read may nevertheless align with the graph that has integrated all the genomes the read does not satisfactorily align with. A situation like this indicates that the read aligns with a hitherto unknown strain of the species represented by the graph, and means that the read stems from that species, which one may overlook when operating with linear genomes alone.

Further, additional advantages of pangenome graphs are due to the compact representation of related genomes. First, the compactness of the graphs renders the execution of alignments simply more efficient: instead of having to align a read with each of the linear genomes separately, one read-to-graph alignment suffices. Secondly, read-to-graph mappings are not only less ambiguous, but also quite simply more accurate, in particular in the presence of structural variations (Garrison *et al*., 2018; Sirén *et al*., 2021). Last but not least, variation graphs, as the (arguably most popular) type of pangenome graphs in use here, facilitate the formulation of an optimization problem that we have termed “path abundance optimization problem”, which gives rise to strain-level taxonomic classification in a tractable way. This optimization problem-based strategy allows for utmost nuanced and accurate distinction between the strains of a species. In summary, read-to-pangenome-graph alignments do not only offer a unified approach to align a read with many related sequences, as mentioned above, but they also offer a way that is unified with respect to the particular type of reads that one deals with. They can be reliably applied for both accurate short and noisy long reads (as well as reads that are both accurate and long, of course), which k-mer-based alignment-free strategies cannot guarantee.

Strain-level classification poses a great methodical challenge, because sequential differences between strains can look near-negligible at first glance. So, strain-level analyses of metagenomes have only recently gained pace. This explains why there are only few existing approaches and why most of them are very recent. PanTax outperforms all other approaches that aim at classifying metagenomes at strain level quite significantly. PanTax does so for both NGS and TGS data, across the entire board of evaluation metrics: It is the only approach that outperforms others significantly while preserving competitive performance rates also in the categories where other approaches have their particular advantages. Last but not least, PanTax proves to outperform the other approaches in strain-level classification of isolate species; although isolates are not the main intention of our approach, these experiments demonstrate that PanTax is not negatively affected by scenarios characterized by a few dominant species, on the contrary.

The main drawback of the original PanTax is its greater demand of computational resources, both in terms of time and memory, compared to other methods, which makes it challenging to process very large databases, such as GTDB. The key steps involved are the construction of pangenome graphs and the loading of them into memory. To address this limitation, we have also developed a fast version of PanTax by incorporating an ultra-fast genome querying step before constructing the pangenome graphs. This approach drastically reduces the number of genomes in the reference database from which the graph is to be constructed, which significantly decreases the time and the memory needed to construct and load the pangenome graphs into main memory. Consequently, PanTax, in its fast version, requires comparable or even fewer computational resources than other approaches. Benchmarking experiments demonstrate that the fast version of PanTax approaches the taxonomic profiling performance rates of the full, default version of PanTax across the full board of datasets; in fact, it even exhibits slight advantages in some cases. Additionally, the accelerated (fast) version of PanTax (fast) handles very large databases, such as GTDB, efficiently and outperforms other methods in terms of taxonomic profiling performance. In summary, PanTax is compatible with NGS, both noisy PacBio CLR and accurate PacBio HiFi, as well as ONT, both noisy R9.4 and the more accurate R10.4 (error rate about 2%) sequencing data. It supports classification at the strain level (and, of course, if desired, also at the level of species), for both single and multiple species, so establishes an approach that is universal with respect to all current scenarios of interest in metagenomics type experiments. While Centrifuge and Centrifuger are the only currently existing methods that rival PanTax in terms of universality, PanTax outperforms Centrifuge/Centrifuger on the vast majority of benchmark categories quite substantially and thus proves to be the preferable universal classifier. We recall that, unlike PanTax, both Centrifuge and Centrifuger are not based on pangenome graphs. This underscores the superiority of pangenome graph based taxonomic classification another time.

Further improvements of PanTax are conceivable, and they are promising. Currently, just like any other reference based classification approach, PanTax is unable to generate taxonomic profiles for strains that because they are novel, are missing in the reference databases. Instead, PanTax reports the most similar strain recorded in the databases. Although this at least accurately determines the most likely closest relative, a preferable solution would be to dynamically update the species-specific pangenome graphs by incorporating hitherto unobserved genetic variation. From a larger perspective, this would support the detection of unknown strains, beyond just enhancing classification by removing mistaken hits.

## Methods

In the following, we provide the full range of methodical details involved in the steps in Figure 1. We continue with additional, relevant remarks regarding benchmark competitors, and we conclude by defining the metrics used for evaluating results.

### Pangenome graph based reference database construction

When constructing our default pangenome based reference database (“Graph representation of reference databases” in Fig. 1), we focus on bacteria as the primary object of interest in the majority of metagenomics studies. We are aware that in metagenomics studies also viruses, archaea, fungi, or plasmids (and so on) can be in the major focus of attention. To account for this, one can, mutatis mutandis, readily extend our methodology to other organisms. Users can flexibly feed their customized databases as input to PanTax, instead of using the provided default databases. Once databases are provided, everything else follows the identical workflow.

In the following, numbers and names of ’Steps’ correspond to the numbering and naming of procedures in Figure 1.

**Step 1: Select high-quality genomes for species.** We retrieved high-quality, complete (in particular gap-free) bacterial genomes from NCBI’s RefSeq (O’Leary *et al*., 2016) on July 18, 2023 (RefSeq Release 219). The NCBI’s RefSeq (release 219) establishes a comprehensive reference database, consisting of 34,697 complete genomes. In view of plasmid sequences being typically considerably shorter than bacterial genomes, plasmid sequences were excluded from our experiments. We remind that one can put particular attention also to plasmids, if desirable, by augmenting our database with plasmid reference sequence based pangenomes, for example. For each species, we computed a pairwise distance matrix using FastANI (Jain *et al*., 2018) where rows and columns correspond to the strains of the species under consideration, and single entries in the matrix indicate the ANI of two strains of that species. Subsequently, we applied a customized graph-based clustering approach (see Algorithm 1 in the Supplementary Methods) at ANI thresholds of 95% and 99.9%, ensuring that the mutual ANI’s of the strain-specific genomes for each species fell within this range. To maintain a balance between sensitivity and computational cost, we determined no more than 10 representative genomes for each species as a reasonable choice. Following these guidelines, we included 8,778 species, amounting to a total of 13,404 strains, in the refined reference database (labeled as RefSeq:13404). Of the 8,778 species included, 1,313 comprised multiple strains (see Supplementary Figure 3). In addition, We used genome updater (https://github.com/pirovc/genome updater) to download genomes from the Genome Taxonomy Database (GTDB). To account for resource limitations during the execution of taxonomic classification tools, we restricted the download to a maximum of 100 genomes per species, resulting in a total of 343,362 strains. The resulting subset of the GTDB genomes was used as our reference Genome Taxonomy Database. We adopt a similar strategy to select high-quality genomes from the Genome Taxonomy Database (GTDB, Release R220) and ultimately construct a refined reference database comprising 206,273 strains (labeled as GTDB:206273).

To handle very large databases, such as GTDB, we developed a fast version of PanTax. The only difference between PanTax (fast) and the original PanTax is the inclusion of an additional step, “ultrafast genome querying” (see Fig. 1). This step is based on the genome querying approach implemented in Sylph (Shaw and Yu, 2024), which utilizes k-mer sketching techniques and a k-mer statistical model to achieve ultrafast and accurate species-level querying. Instead of constructing pangenome graphs using all genomes in the reference database, PanTax (fast) first identifies the species potentially present in the sequencing data using this highly efficient strategy. This enables quick filtering at the species level, thereby reducing the number of pangenome graphs that need to be constructed.

**Step 2: Construct a pangenome graph for each species.** Subsequently, we constructed a variation graph (as the particular, and generally most popular type of pangenome graph) for each species, by using the (efficient and unbiased) pangenome graph builder PGGB (Garrison *et al*., 2023). We construct the variation graph *G* = (*V, E, P* ) from the (multiple) strain genomes pertaining to a species, where *V* are the nodes, *E* are the edges and *P* are the paths in *G* corresponding to the input strain-specific genomes used for constructing the graph. *G* is a bidirected sequence graph, where each vertex *v* ∈ *V* corresponds to a sequence seq(*v*) that consists of single or multiple nucleotides (A, T, C, G).

Edges *e_ij_* = (*v_i_, v_j_*) ∈ *E* indicate that the sequences seq(*v_i_*) and seq(*v_j_*) represented by nodes *v_i_*and *v_j_*appear as a sequential subsequence in one of the genomes that contribute to the graph. Because paths *P* outline these genomes, paths *P* cover the graph, or, in other words, each edge is contained in one of the paths from *P* . Let 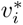 denote the reverse-complement of node *v_i_*. Then the reverse edge 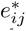 of *e_ij_* is defined as 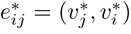. Both nodes and edges can be traversed in either the forward or the reverse direction in the graph. In general, known strains can be identified as the specific paths *p* = (*v*_1_*, …, v_l_*) ∈ *P* , as part of the definition of the species-specific pangenome graph *G*.

For species that involve only one strain, the task is to turn its corresponding sequence into a dedicated variation graph, acting as representative counterpart to the pangenome graphs constructed from multiple strains. Here, we achieved this by partitioning the genome sequence into fragments of (the relatively short) length 1024bp, each of which is to correspond to a node in the graph, which, by design, is chain-like.

Subsequently, one merges all species-specific pangenome graphs into a single, large pangenome graph, by relabeling nodes, while never introducing new nodes or edges. Of note, the species-specific pangenome graphs can be computed in paraellel, which avoids computational bottlenecks in this step. As outlined above, this procedure supports the unified and unambiguous classification of single reads via *one* read-to-graph alignment, instead of having to align a read with each of the (here, by default: 8,778) species-specific pangenome graphs individually. In other words, instead of having to perform and evaluate 8,778 alignments in a statistically involved mutual context, the merging of graphs delivers one encompassing, optimal and statistically sensible alignment that directly indicates the species the read optimally aligns with.

### Species-level taxonomic classification

**Step 3: Sequence to graph alignment.** Sequencing reads are then mapped to the merged pangenome graph using existing, approved and efficient sequence-to-graph aligners. For short reads, we employ Giraffe (Sirén *et al*., 2021), as an integral part of the vg toolkit (Garrison *et al*., 2018) supporting the computation of sequence-to-graph alignments for preferably short reads. For long reads, we make use of GraphAligner (Rautiainen and Marschall, 2020) by default.

For faster alignment of long reads, we also have implemented an option (--fast-aln) that utilizes the Giraffe aligner in its long-read mode instead of GraphAligner. Note that the long read mode of Giraffe is currently under development such that alignments tend to be worse than those computed with GraphAligner. Despite the current disadvantages of Giraffe when using long reads, we have nevertheless implemented it, in expectation of future progress thanks to the rapid developments it is currently undergoing.

The basis for speeding up read alignment with Giraffe is the construction of a graph Burrows-Wheeler transform (GBWT) (Sirén *et al*., 2020) which one uses for indexing the pangenome graph. For PanTax, we resort to building the GBWT index manually, via a ’fast indexing mode’, which relies on three commands provided by vg, which considerably speeds up index construction. Although the default fully automatic construction of the index via ’vg autoindex’ is supposed to be more accurate, classification results did not differ in our experiments, which explains our choice.

**Step 4: Species-level taxonomic binning.** Taxonomic binning involves the allocation of sequencing reads to distinct taxonomic groups. In this procedure, we assign each mapped read to a species-specific pangenome graph, utilizing the results of the sequence-to-graph alignment resulting from aligning reads with the merged pangenome graph. When a read aligns to multiple graphs (within this unifying merged graph), we only retain the optimal alignment, thereby ensuring that each read corresponds to a single species. We again note that the optimal alignment is easy to determine precisely because the merging of graphs implies that different alignment scores can be put into mutual context.

Subsequently, we assess whether individual species need to be flagged as false positives. That is, although the species receives a high score, the species does not make part of the metagenome. For that, we examine the MAPQ (mapping quality) scores of all the reads whose optimal alignments correspond to the particular species. We raise two key criteria: first, we require at least one read to reach a MAPQ of 60, which indicates a statistically significant unique match of at least one read with that species. We further require that at least one tenth of the reads that optimally align with a species exhibit a MAPQ of at least 2; if less than one tenth of the reads aligned with a species have MAPQ less than 2, the species is flagged as false positive. It remains to mention that reads may fail to align with the pangenome graph, which potentially indicates the presence of novel strains (not necessarily from novel species, although that is possible as well) in the metagenomic sample under analysis. We discard such unmapped reads from further consideration in the subsequent analysis.

**Step 5: Species-level taxonomic profiling (optional)** For species *i*, we divide the total base count of the reads that optimally aligned with species *i* via the read-to-graph alignment by the average length of the strain-specific genomes that contributed to constructing the pangenome graph for species *i*, which we refer to as *normalized read coverage c_i_* in the following. Subsequently, we calculate the relative (taxon) abundance of species *i* as the ratio *c_i_/ _i_*_∈_*_T_ c_i_*, where *T* is the set of species reported in the previous step (e.g. resulting from excluding potentially false positives when dealing with NGS data, before outputting all species referred to as *T* here). Note that this step is not necessary for subsequent strain-level classification; it is optional unless explicitly required by the user.

### Strain-level taxonomic classification

**Step 6: Path abundance optimization.** Furthermore, we offer strain-level classification, reflecting the main purpose of our approach. Having already constructed a species-specific pangenome graph by collapsing identical sequences from multiple strains (as described in Step 2), we formulate a path abundance optimization problem for each species-specific pangenome graph, following an approach suggested in (Baaijens *et al*., 2020). In this step, the abundance we refer to is absolute abundance, which is the average coverage depth. This step aims to resolve strain-level taxonomic classification. Let *V* be the set of all nodes in a species-specific pangenome graph, *P* be the set of paths corresponding to all strains used to construct the pangenome graph for a particular species (where a strain’s genome sequence corresponds to a path in the pangenome graph in general), and *a_v_* be the absolute abundance of node *v* ∈ *V* in the graph, as estimated by evaluating the reads whose alignments include *v* (i.e., the average coverage depth of a single base in this node), where we follow the definitions provided in (Baaijens *et al*., 2020) in a one-to-one fashion. For a path *p* ∈ *P* (again following (Baaijens *et al*., 2020)), we define *a_p_* ∈ R_≥0_ to be the absolute abundance of path *p*, which corresponds to the abundance of the strain that corresponds to *p*. Here, we would like to predict all *a_p_* by raising them as variables in the program

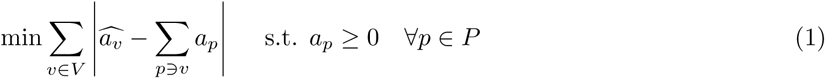

which we refer to as *path abundance optimization (PAO)* problem. Values *a_p_* ≥ 0 establishing a solution are output as the abundance values of the strains that correspond to the paths *p*.

As for an explanation, the objective function of the PAO problem aims to estimate the abundance (i.e. the normalized read coverage) of each strain by minimizing the difference between the abundances of the nodes one has observed (derived from the sequence-to-graph alignments in Step 3) and the abundances of the nodes that correspond to the abundances of the strains whose genomes include the nodes. The PAO problem is convex in nature, and can be efficiently linearized and solved using a linear programming solver of choice; here we opted for Gurobi (Gurobi Optimization, 2023). So, from a theoretical perspective, this problem is polynomial-time solvable. In addition, since the size of the candidate set of strain paths *P* is typically not overly large (usually ≤ 10), the optimization problems can be solved in parallel across the species, which renders this step computationally very efficient.

Note that for each species, we conduct two iterations of path abundance optimization. To further mitigate the odds of raising false positives during classification, we add a filtering operation prior to each iteration of PAO. To illustrate the filtering step, we first define a *triplet node* as a set of three consecutive nodes within a path. A triplet node that is exclusive to a single strain, that is the three nodes only appear in that order in one of the paths *p* ∈ *P* , is referred to as *strain-specific triplet node*.

1. For the filtering step that precedes the first iteration of solving the PAO, let *V*_reads_ be the consecutive node triples covered by reads aligning with the species-specific pangenome graph and *V*_strain_ reflect the set of strain-specific triplet nodes that show in the path corresponding to the strain, as just defined. We then raise the metric f, as per the definition

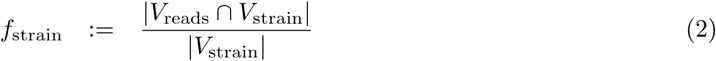

As per its definition, *f* captures the proportion of strain-specific triplet nodes in the strain genome (reference) the concatenation of the three sequences of which appears in the reads of the sequencing sample. As a ratio, the value of *f* naturally ranges from 0 to 1, with 1 indicating that all strain-specific triplet nodes are covered by reads from the sample.

Let **v**_triplet_ refer to strain-specific triplet nodes, as defined above, and let *c*(**v**_triplet_) be the average read coverage of **v**_triplet_, where averaging is across nucleotides in the sequence of **v**_triplet_. Further, we introduce *a_p,_*_triplet_, which, unlike *a_p_* that resulted from the solution of the PAO, estimates the abundance of the strain that corresponds to *p* by taking the average of the coverages of the triplet nodes, that is the average of the *c*(**v**_triplet_) in *p*:

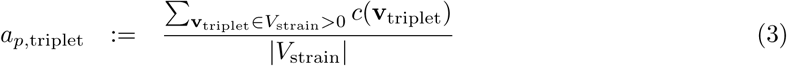

Subsequently, we solve the PAO problem a first time for only the strains that were not discarded in the filtering procedure (that is whose *f*_strain_ was found to be ≥ 0.3 for broad multi-species tasks or ≥ 0.43 for single-species tasks), and record the strains, respectively the paths *p* in the species-specific pangenome graph to which they correspond whose *a_p_ >* 0, that is we keep all strains that make part of the solution of the PAO.

1. For the filtering step that precedes the second iteration of solving the PAO, we raise the metric

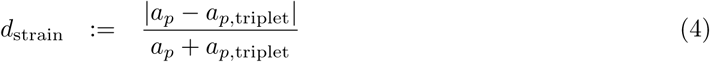

As for a brief explanation, *d*_strain_ quantifies the divergence between the strain abundance *a_p_* predicted in the first iteration of the PAO and the strain abundance estimate based on the average coverage of all strain-specific triplet nodes.

Before proceeding with the second iteration of the PAO, we filter out further potentially false positive strains by keeping only strains where *d*_strain_ ≤ 0.46. At this stage, for strains with 0.46 *< d*_strain_ ≤ 0.6, we perform a rescue operation for high-confidence strains among them. We define the base coverage of a strain as *b*_strain_. A strain will be retained if it satisfies the condition *b*_strain_ ∗ *f*_strain_ ≥ 0.85.

Finally, we solve the PAO a second time. The final output of strains are the ones whose *a_p_* is greater than zero in that second iteration. For the rescued strains, the minimum value between the PAO final solution *a_p_* and *a_p,_*_triplet_ is selected as their absolute abundance.

**Step 7: Strain-level taxonomic profiling.** We firstly defined a metric *d*_species_ for filtering, which quantifies the divergence between the absolute abundance *c_i_* of a species *i* (as described in Step 5) and the total absolute abundance of all its strains *c_i,strains_*as determined by the PAO solution

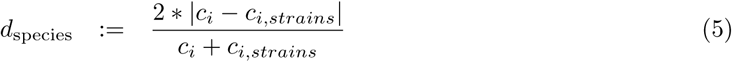

We filter out further potentially false positive species and its strains with *d*_species_0.2. The divergence is typically caused by reads originating from one species being aligned to another non-existent species.

These reads often come from shared regions, leading to their excessive accumulation in these areas. Our objective is to filter out these non-existent species.

The final, additional step is to provide relative abundance estimates for any strain predicted to make part of the sample. For that, one determines the absolute abundances of the strains, and divides them by the absolute abundance of the species they belong to, as predicted in step 5. There is one caveat: the sum of the absolute abundances of the strains could exceed the absolute abundance of the most abundant strain within the species. In that case, we scale the absolute abundances of the strains such that their sum equals that of their species. The relative abundance of one strain then is the appropriately normalized / scaled version of the absolute abundance of the strain, as just determined.

### Metrics for evaluation

We evaluate the performance of taxonomic classifiers using metrics that are widely adopted in various relevant studies (Simon *et al*., 2019; Zhu *et al*., 2022; Luo *et al*., 2022a). The primary criteria for assessing metagenomic classification methods are precision and recall. At the level of taxa (strain level that we focus on), precision measures the fraction of correctly identified taxa among all taxa that were identified. Recall, at the level of taxa, evaluates the proportion of correctly identified taxa compared to all taxa present in the sample. The F1 score, the harmonic mean of precision and recall, is frequently employed as a balanced metric for evaluating classifiers.

However, precision, recall, and the F1 score may not reflect the performance of classifiers in metagenomic samples sufficiently comprehensively, because the F1 score neglects the correct assessment of low-abundance taxa, whose accurate detection, however, can be crucial. A approach to evaluating the contents of metagenomes that takes low-abundance taxa into sufficiently accurate account is to evaluate the metagenomes in terms of curves that plot precision and recall across varying abundance thresholds. Calculating the area under this curve (“AUPR”) yields a metric that does not neglect the performance of a classifier also with respect to taxa of lower abundance (Simon *et al*., 2019). For the calculation of AUPR, refer to (Simon *et al*., 2019). Classifiers that fail to recall all ground-truth taxa are penalized with a zero AUPR score from their greatest recall achieved up to 100% recall. Conversely, classifiers that achieve 100% recall are not further penalized in their AUPR score for additional false positive taxon calls. Please see “AUPR calculation example” in the Supplementary Methods for illustrative examples.

Beyond the accurate identification of taxa only, it can be imperative to provide accurate estimates of the abundances of the taxa that make part of a metagenome. An example for the necessity of such practice are metagenome-wide association studies (“MetaGWAS”) where fluctuations in the proportions of the participating taxa can have profound implications in terms of the phenotypes supported by the metagenome. The relative abundances of taxa across various samples are comparable, making them a primary focus in MetaGWAS studies. The ultimate output of taxonomic profiling at the strain level (Steps 7) is precisely these relative abundances. Notably, in this study, we propose a set of metrics to assess the accuracy and reliability of these relative abundance estimates. To determine abundance profiles for tools that can output read assignment, we follow the exact same protocol employed for PanTax. Namely, all tools use relative taxonomic abundance for evaluation, where the genome length of a strain is used to normalize read coverage. If methods offer taxonomic abundance profiling as output directly, we make use of that. The L1 distance, L2 distance and BC distance, commonly used metrics, serve as a quantitative evaluation of abundance profiles (Sun *et al*., 2021; Simon *et al*., 2019). Given a sample, let *I* represent the intersection set of estimated and true taxa. For a taxon *i* ∈ *I*, let 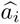 and *a_i_* represent the estimated and true abundance of taxon *i*, respectively. Then, the L1 distance is computed as follows: 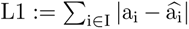. The L2 distance is computed as follows: 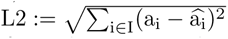. The calculation of the BC distance is performed using the *braycurtis* function from the *scipy* Python package, and it is defined as 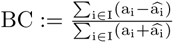. A smaller distance indicates a higher similarity between the estimated and true abundances, while greater distance suggests greater divergence. However, it is worth noting that abundance profile distance is particularly sensitive to the accurate quantification of highly abundant taxa within a sample, as these taxa exert a significant influence on the overall similarity between abundance profiles. To address this, we introduce two complementary metrics: the absolute frequency error (AFE) and the relative frequency error (RFE). AFE is defined as: 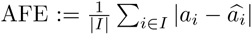, whereas RFE is defined as: 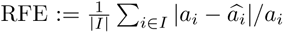. These two metrics have been adapted from previous studies on viral haplotype abundance evaluation (Luo *et al*., 2022a; Baaijens *et al*., 2020). The AFE and RFE metrics provide insights into the deviation of estimated abundances from their true values. Notably, the RFE metric balances the influence of low-abundance species or strains, because it scales divergences relative to the true abundance *a_i_*of the taxa.

## Data availability

The genomes and simulated sequencing reads generated by this study can be downloaded from Zenodo DOI: https://doi.org/10.5281/zenodo.15148106.

## Code availability

The source code of PanTax is GPL-3.0 licensed, and publicly available at https://github.com/ LuoGroup2023/PanTax. The results presented in this study can be reproduced from Code Ocean under DOI: https://codeocean.com/capsule/6598572/tree/v1.

## Acknowledgements

XL is supported by the National Natural Science Foundation of China (Grant No. 32400506), the Natural Science Foundation of Hunan Province (Grant No. 2024JJ4008) and Fundamental Research Funds for the Central Universities (Grant No. 541109030062). AS received funding from the European Union’s Horizon 2020 research and innovation programme under Marie Sk-lodowska-Curie grant agreements No 956229 (ALPACA) and No 872539 (PANGAIA). YL is supported by the National Natural Science Foundation of China (Grant No. 62372159).

## Author contributions

Xiao Luo conceived this study. Xiao Luo, Wenhai Zhang and Yuansheng Liu designed the method. Wenhai Zhang implemented the software. Xiao Luo and Alexander Schönhuth wrote and revised the manuscript. Wenhai Zhang, Yuansheng Liu, Guangyi Li, Jialu Xu and Enlian Chen conducted the data analysis. All authors read and approved the final version of the manuscript.

